# MK2-deficient mice are bradycardic and display delayed hypertrophic remodelling in response to a chronic increase in afterload

**DOI:** 10.1101/2020.01.23.916049

**Authors:** Matthieu Ruiz, Maya Khairallah, Dharmendra Dingar, George Vaniotis, Ramzi J. Khairallah, Benjamin Lauzier, Simon Thibault, Joëlle Trépanier, Yanfen Shi, Annie Douillette, Bahira Hussein, Sherin Ali Nawaito, Pramod Sahadevan, Albert Nguyen, Marc-Antoine Gillis, Martin G. Sirois, Matthias Gaestel, William C. Stanley, Céline Fiset, Jean-Claude Tardif, Bruce G. Allen

**Affiliations:** Department of Medicine, Université de Montréal, Montréal, Québec, Canada, H3C 3J7; Department of Biochemistry and Molecular Medicine, Université de Montréal, Montréal, Québec, Canada, H3C 3J7; Department of Pharmacology and Physiology, Université de Montréal, Montréal, Québec, Canada, H3C 3J7; Department of Faculté de Pharmacie, Université de Montréal, Montréal, Québec, Canada, H3C 3J7; Montreal Heart Institute, 5000 Belanger St., Montréal, Québec, Canada, H1T 1C8; University of Maryland, Baltimore, MD, 212101, USA; L’Institut du thorax, INSERM, CNRS, Université de Nantes, Nantes, France; Institute of Cell Biochemistry, Hannover Medical School, Carl-Neuberg-Strasse 1, 30625 Hannover, Germany; Department of Physiology, Faculty of Medicine, Suez Canal University, Ismailia, Egypt

**Keywords:** p38 MAPK, MK2, bradycardia, cardiac remodeling, mitochondrial permeability transition pore

## Abstract

MAP kinase-activated protein kinase-2 (MK2) is protein serine/threonine kinase activated by p38α/β. Herein we examined the cardiac phenotype of pan MK2-null (MK2^−/−^) mice. Survival curves for male MK2^+/+^ and MK2^−/−^ mice did not differ (Mantel-Cox test, *P* = 0.580). At 12-weeks of age, MK2^−/−^ mice exhibited normal systolic function along with signs of possible early diastolic dysfunction; however, ageing was not associated with an abnormal reduction in diastolic function. Both R-R interval and P-R segment durations were prolonged in MK2-deficient mice. However, heart rates normalized when isolated hearts were perfused *ex vivo* in working mode. Ca^2+^ transients evoked by field stimulation or caffeine were similar in ventricular myocytes from MK2^+/+^ and MK2^−/−^ mice. MK2^−/−^ mice had lower body temperature and an age-dependent reduction in body weight. mRNA levels of key metabolic genes, including *Ppargc1a*, *Acadm*, *Lipe*, and *Ucp3* were increased in hearts from MK2^−/−^ mice. For equivalent respiration rates, mitochondria from MK2^−/−^ hearts showed a significant decrease in Ca^2+^-sensitivity to mitochondrial permeability transition pore (mPTP) opening. Finally, the pressure overload-induced increase in heart weight/tibia length and decrease in systolic function were attenuated in MK2^−/−^ mice two weeks, but not eight weeks, after constriction of the transverse aorta. Collectively, these results implicate MK2 in (i) autonomic regulation of heart rate, (ii) cardiac mitochondrial function, and (iii) the early stages of myocardial remodeling in response to chronic pressure overload.

**Key points summary:** The cardiac characterization of pan MK2-null mice showed:

i. altered autonomic regulation of heart rate
ii. increased expression of key metabolic genes
iii. decreased Ca^2+^-sensitivity for MPTP opening
iv. delayed hypertrophic remodeling in response to increased afterload

## Introduction

The primary response of the heart to increased workload, especially in reaction to pressure or volume overload, is to adapt myocyte size in an attempt to reduce ventricular wall stress and/or compensate for increased hemodynamic demand (Frey *et al.*, 2004). However, unlike physiological hypertrophy, in response to exercise or during post-natal growth, stress stimuli such as arterial hypertension or myocardial infarction (Frey *et al.*, 2004) leads to pathological hypertrophy or hypertrophic cardiomyopathy, associated with interstitial fibrosis and re-expression of a fetal gene program, which can result in increased myocardial stiffness and reduced cardiac output (Sadoshima & Izumo, 1997; Depre *et al.*, 1998; Kong *et al.*, 2014). Although cardiac hypertrophy is seen as a compensatory phenomenon to normalize myocardial wall stress, when sustained, cardiac hypertrophy predisposes to sudden cardiac death, arrhythmia, and development of heart failure (Frey *et al.*, 2011). Hence, understanding the basic signaling pathways involved in maintaining optimal cardiac function and their role in pressure overload-induced hypertrophy, as well as the associated contractile dysfunction, is necessary for the development of more effective means of prevention and treatment.

Several signaling pathways have been implicated in the transition from compensated hypertrophy to decompensated heart failure, including the mitogen-activated protein kinase (MAPK) pathway (Shah & Mann, 2011) in which p38 MAPK plays a central role (Trempolec *et al.*, 2013a). Increased activation of p38 MAPK has been observed both in experimental and human heart failure (Takeishi *et al.*, 2002; Bellahcene *et al.*, 2006; Arabacilar & Marber, 2015; Chahine *et al.*, 2015). However, the pathological cardiac phenotype associated with p38 MAPK signaling is, to some extent, confusing. The literature contains seemingly contradictory reports indicating p38 activation is both beneficial and detrimental during hypertrophy. Four isoforms of p38 MAPK (p38α, p38β, p38γ, and p38δ) are expressed in the heart (Lemke *et al.*, 2001; Dingar *et al.*, 2010): p38α and p38β have been shown to be activated during pressure overload and this activation correlates with ventricular hypertrophy (Wang *et al.*, 1998; Yokota & Wang, 2016). Similarly, p38γ and p38δ have been shown to mediate hypertrophy in response to angiotensin II infusion (Gonzalez-Teran *et al.*, 2016). Furthermore, in isolated cardiomyocytes, activation of p38 using either pharmacological or genetic approaches also suggest p38 activation is pro-hypertrophic (Nemoto *et al.*, 1998; Wang *et al.*, 1998). The latter is reinforced by the attenuation of the hypertrophic response, evoked with phenylephrine or endothelin-1, using inhibitors of p38 (Nemoto *et al.*, 1998). In contrast, chronic inhibition of p38 signaling using dominant-negative p38α or β is associated with enhanced cardiac hypertrophy in response to pressure overload or following infusion with angiotensin II, isoproterenol, or phenylephrine (Braz *et al.*, 2003; Zhang *et al.*, 2003). Similarly, cardiomyocyte-targeted deletion of p38α in mice did not impede hypertrophy in response to aortic banding, but increased both myocyte apoptosis and fibrosis while decreasing in cardiac contractility (Nishida *et al.*, 2004). In contrast, myocyte-targeted activation of p38α in adult mice induced hypertrophy, fibrosis, and mortality within one week (Streicher *et al.*, 2010).

Although there are divergent results, all convey a similar message, in which p38 MAPK signaling is essential for heart function and plays an important role in the pathogenesis of heart failure. However the apparent opposing and divergent role for p38, as well as its numerous targets/substrates that are involved in a wide variety of cellular processes (Cargnello & Roux, 2011; Trempolec *et al.*, 2013b), makes it difficult to predict the impact of p38 inhibition in clinical trials. Moreover, none of the p38 inhibitors developed to date translated into safe and effective clinical strategies with low and acceptable toxicity due to their unwanted systemic side effects, which include hepatotoxicity, neurotoxicity, and/or cardiotoxicity (Coulthard *et al.*, 2009; Marber *et al.*, 2011; Martin *et al.*, 2015). Hence, further research to better understand the role of p38 MAPK signaling in the healthy heart, especially with regards to its downstream targets, is essential to elucidate the potential therapeutic implications of targeting this signaling pathway. Over the last decade, many studies have focused on MK2, among other downstream p38 targets. Several reports suggested that targeting MK2 could lead to beneficial effects equivalent to those obtained when directly inhibiting p38 but with the additional advantage of lacking the side effects associated to p38 inhibitors (Fiore *et al.*, 2016) and could therefore serve as a potential therapeutic target. Indeed, the side effects associated with p38 inhibitors were recently attributed to the loss of feedback control of upstream TAB1/TAK1 signaling, which in turn leads to sustained activation of pro-inflammatory pathways (Cheung *et al.*, 2003; Arthur & Ley, 2013). In contrast, this feedback control is maintained when using MK2 inhibitors, thereby preventing collateral activation of pro-inflammatory pathways (Dulos *et al.*, 2013). In addition, evidence suggests that MK2 may be an interesting target in other forms of cardiomyopathy (Ruiz *et al.*, 2018). In fact, MK2 deficiency protects against ischemia-reperfusion injury (Shiroto *et al.*, 2005). Streicher *et al*. also reported that deleting MK2 in a conditional and cardiomyocyte-specific mouse model expressing constitutively active MKK3bE significantly reduces the MKK3bE-induced hypertrophy, improves contractile performance and rescued lethality (Streicher *et al.*, 2010). In addition, pharmacological inhibition of MK2 reduces cardiac fibrosis and preserves cardiac function following myocardial infarction (Xu *et al.*, 2014). Finally, we have recently shown that the absence of MK2 prevents diabetes-induced cardiac dysfunction and perturbations in lipid metabolism (Ruiz *et al.*, 2016; Ruiz *et al.*, 2018).

To date, few studies have examined the role of MK2 in either the healthy heart or in myocardial remodeling in response to a chronic increase in cardiac afterload. Therefore, using a previously-described MK2-deficient mouse model (Kotlyarov *et al.*, 1999), this study was undertaken to investigate (i) whether MK2 is involved in maintaining basal cardiac function and (ii) whether MK2 plays a role in cardiac remodeling resulting from chronic pressure overload, in which case it would represent a potential therapeutic target.

## Methods

### Ethical approval

All animal experiments were approved by the local ethics committee and performed according to the guidelines of the Canadian Council on Animal Care.

### Animals

Twelve-week-old male wild-type (MK2^+/+^) and MK2-deficient (MK2^−/−^) littermate mice were used. These mice have been previously described (Kotlyarov *et al.*, 1999). MK2^−/−^ mice are viable, fertile and show no behavioral or physiological defects.

### *In vivo* cardiac function

#### Radio telemetry

Radio telemetry was used for continuous recordings of heart rate and ECG data in conscious unrestrained mice. MK2^+/+^ mice and age-matched MK2^−/−^ mice were instrumented, under isoflurane anaesthesia (2.5% in O_2_, 0.5 l/min for induction and 1.5% for maintenance), with OpenHeart single-channel telemetry transmitters (Data Sciences International, Arden Hills, MN, USA) as described previously (Brouillette *et al.*, 2007). For analgesia, buprenorphine (0.05 mg/kg) was administered before and every 8 h for 48 h after surgery. Leads were placed in the conventional electrocardiogram lead II position with the positive transmitter lead located on the left anterior chest wall above the apex of the heart and the negative lead located on the right shoulder. Mice were provided with food and water ad libitum and maintained on a 12:12 h day-night cycle. One-week post-surgery, data acquisition was initiated at a sampling frequency of 1 kHz and maintained for 48 h. Recordings were analysed with ECG Auto (version 2.8; EMKA Technologies, Paris, France). Briefly, the first 10 minutes of every 3-hour period was analyzed for the entire 48 hours of recording. A smoothing of 1 ms and notch filter of 60 Hz were used to reduce noise. Signals were averaged from 10 consecutive beats. ECG parameters were obtained through an automated signal detection using typical traces. The analyst was blinded as to genotype and the mice shared the same library of typical traces. The QT interval was corrected for differences in R-R interval using the standard formula for mice (QTc=QT/(RR/100)^1/2^) (Mitchell *et al.*, 1998). Heart rate variability was analyzed using Kubios HRV Standard (Version 3.0.0) to fit an ellipse to a Poincaré plot (R-R_n_ versus R-R_n+1_) of R-R interval data. Results were expressed as the ratio of the standard deviation on the short axis (SD1) versus the standard deviation on the long axis (SD2) of the ellipse. SD1 reflects short term HRV whereas SD2 reflects both short term and long term HRV (Shaffer & Ginsberg, 2017).

#### Surface electrocardiograms

Surface ECG recordings were assessed on male MK2^+/+^ and MK2^−/−^ mice, under anesthesia with 2% isoflurane (100% O_2_. flow rate of 1 L/min), before and after intravenous (IV) injection of isoproterenol (0.1 μg/g) as previously described (El Khoury *et al.*, 2013).

#### Transthoracic echocardiography

Transthoracic echocardiographic imaging was performed on mice, sedated with isoflurane (2% in 100% O_2_), 1 d before constriction of the transverse aorta (TAC), for baseline evaluation, and 1 d before sacrifice. In the 8-wk cohort, echocardiographic imaging was also performed 2-wk post-surgery. Examinations were carried out with a linear array i13L probe (10-14 MHz) using a Vivid 7 Dimension ultrasound system (GE Healthcare Ultrasound, Horten, Norway). The operator was blinded to the genotype of the mice. All parameters and calculations related to left ventricular (LV) and right ventricular (RV) structure, LV and RV systolic function, LV and RV diastolic function, left atrium dimensions, and myocardial performance index were obtained as previously described and are briefly described in online supplemental information (Merlet *et al.*, 2016; Nawaito *et al.*, 2017).

#### Millar catheterization

Hemodynamic parameters were evaluated using a Mikro-Tip pressure transducer catheter (Millar instruments) in mice anesthetized with 2% isoflurane (100% O_2_. flow rate of 1 L/min). Body temperature was monitored and maintained at 37°C using a heating pad. The catheter was inserted into the left ventricle through the carotid as previously reported (Nawaito *et al.*, 2017). Recordings were analyzed using IOX software version 2.5.1.6 (EMKA technologies).

#### Transverse aortic constriction

TAC was performed in 12-wk-old male mice sedated with isoflurane gas as previously described (Nawaito *et al.*, 2017). The surgeon was blinded as to the genotype of the mice. Sham animals underwent the identical surgical procedure, but the aorta was not constricted. Two or eight wk after surgery, mice were anesthetized with pentobarbital and sacrificed, hearts removed, weighed, snap-frozen in liquid nitrogen-chilled 2-methyl butane, and stored at −80 °C. Pentobarbital, rather than isoflurane, was always used prior to sacrifice as isoflurane activates p38 MAP kinase in the mouse heart (Data Not Shown).

### *Ex vivo* cardiac function

#### Ex vivo working heart perfused in semi-recirculating mode

Hearts were isolated from fed mice and perfused under normoxic conditions for 30 minutes to evaluate basal function. To assess the impact of isoproterenol, hearts from a second group of mice were perfused under normoxic conditions for 20 minutes after which isoproterenol (10 nM) was added to the buffer and perfusion continued for an additional 20 min. All perfusions were carried out at a fixed preload (15 mmHg) and afterload (50 mmHg, except following addition of isoproterenol (see below), and using a semi-recirculating modified Krebs-Henseleit buffer containing a mix of substrates and hormones as previously described (Khairallah *et al.*, 2004; Ruiz *et al.*, 2015) in the absence (11 mM glucose, 1.5 mM lactate, 0.2 mM pyruvate, 50 mM carnitine, 0.8 nM insulin, and 5 nM epinephrine) or presence of palmitate (11 mM glucose, 1.5 mM lactate, 0.2 mM pyruvate, and 0.7 mM palmitate bound to 3% dialyzed albumin, 50 mM carnitine, 0.8 nM insulin, and 5 nM epinephrine). During perfusion in the presence of isoproterenol (10 nM) the afterload was increased to 60 mmHg. Functional parameters were monitored throughout the perfusion (iox2 data acquisition software, EMKA Technologies): data shown were acquired during the final 5 min of each perfusion.

### *In vitro* analyses on isolated ventricular myocytes or mitochondria

#### Ca^2+^ transients and caffeine-induced Ca^2+^ transients

Ca^2+^ transients and sarcoplasmic reticulum Ca^2+^ content were assessed in cardiac ventricular myocytes isolated from MK2^+/+^ and MK2^−/−^ mice as described previously (Rivard *et al.*, 2011).

#### Isolation of cardiac mitochondria and mitochondrial respiration measurements

Subsarcolemmal mitochondria (SSM) were isolated from adult mouse hearts as described previously (Khairallah *et al.*, 2010b). SSM were resuspended in KME buffer (100 mM KCl, 50 mM MOPS, 0.5 mM EGTA) at a final concentration of 25 mg mitochondrial protein/ml. To assess respiration, SSM were suspended (0.25 mg protein in 0.5 ml) in a respiration buffer containing 100 mM KCl, 50 mM MOPS, 5 mM KH_2_PO_4_, 1 mM EGTA, and 0.1% fatty acid-free BSA, pH 7.0. Oxygen consumption was measured using a Clark-type electrode. After recording the basal respiration rate, the following substrate conditions were assessed: (i) 10 mM glutamate plus 5 mM malate, (ii) 10 mM pyruvate plus 5 mM malate, (iii) 10 mM palmitoyl carnitine plus 5 mM malate, and (iv) 10 mM succinate plus 7.5 µM rotenone as previously described (Khairallah *et al.*, 2010b). State III respiration was measured in the presence of 200 µM ADP and state IV respiration was measured with the addition of oligomycin (4 μM).

#### Assessing Ca^2+^-induced mitochondrial permeability transition pore opening

The calcium-dependency for mitochondrial permeability transition pore (MPTP) opening in SSM was determined as previously described (O’Shea *et al.*, 2009). In short, mitochondria (0.5 mg) were resuspended in 2.0 ml of assay medium containing 100 mM KCl, 50 mM MOPS, 5 mM KH_2_PO_4_, 5 µM EGTA, 1 mM MgCl_2_, 5 mM glutamate, and 5 mM malate and assayed for Ca^2+^ uptake in a fluorescence spectrophotometer at 37°C. CaCl_2_ (5 mM) was infused at a rate of 2 µl/min and the concentration of extramitochondrial free Ca^2+^ was determined using 0.1 mM Fura-6F by monitoring fluorescence emission at 550 nm with excitation wavelengths for the free and calcium-bound forms of 340 and 380 nm, respectively. The cumulative Ca^2+^ load that was required to induce MPTP opening was determined from a semi-log plot of the extramitochondrial [Ca^2+^] versus the cumulative Ca^2+^ load and defined as the amount of infused Ca^2+^ required to induce a sharp increase in extramitochondrial [Ca^2+^] (Khairallah *et al.*, 2010a). Analyses were done by a single investigator who was blinded as to the identity of the samples.

#### Measuring reactive oxygen species

To determine whether abolishing MK2 activity affected mitochondrial generation of reactive oxygen species, hydrogen peroxide production was measured in respiring mitochondria as previously described (O’Shea *et al.*, 2009). Briefly, 0.75 mg of mitochondria were incubated in respiration buffer containing malate and glutamate to which 5 U/ml horseradish peroxidase, 40 U/ml Cu-Zn superoxide dismutase, and 1 µM Amplex Red were added. Superoxide generation was measured with sequential additions of ADP (0.5 mM), oligomycin (1.25 µg/ml) and rotenone (1 µM). H_2_O_2_ production was measured as an increase in fluorescence of Amplex Red. The experiment was terminated by the addition of 1 nmol H_2_O_2_ to calibrate the dye response.

### Cardiac tissue analyses

#### Histological analysis

Hearts were embedded in Tissue-Tek O.C.T. compound (Sakura Finetek USA, Inc.) and transverse cryosections (8 μm) of the ventricles were prepared and stained with Masson trichrome. Images were taken at X40 using an Olympus BX46 microscope. Collagen content was quantified using Image Pro Plus version 7 (Media Cybernetics, USA) and expressed as a percentage of the surface area. Perivascular collagen was excluded from the measurements.

#### Quantitative real time-PCR

Total cellular RNA was isolated from transverse cryosections (14 μm) of murine ventricular myocardium using RNeasy Micro kits (Qiagen Inc.) with minor modifications. Total RNA was extracted by vortexing tissue section in 300 μl TRIzol reagent (Sigma-Aldrich) for 30 seconds. After incubating at ambient temperature for 5 minutes, 60 μl chloroform was added, samples were again vortexed, and maintained at ambient temperature for an additional 2-3 minutes. After centrifugation for 15 minutes at 18,300 x g and 4°C, the upper aqueous phase was collected, diluted with an equal volume of 70% ethanol, and total RNA purified on Qiagen columns according to the manufacturer’s instructions. Finally, total RNA was eluted with 14 μl of distilled, RNAse-free water. cDNA synthesis was performed as described previously (Dingar *et al.*, 2010). qPCR was performed by single observer in a blinded manner. The following genes were selected: (i) key metabolic enzymes, namely medium-chain acyl-CoA dehydrogenase (*Acadm*) and hormone-sensitive lipase (*Lipe*), (ii) a transcriptional coactivator acting as a master regulator of mitochondrial function and regulating metabolic gene expression namely peroxisome proliferator-activated receptor γ coactivator 1-α (*Ppargc1a*), (iii) mitochondrial uncoupling protein 3 (*Ucp3*), (iv) a marker gene of the fetal phenotype, namely myosin heavy chain β (*Myh7*), and (iv) a marker of early cardiac remodeling, namely atrial natriuretic factor (*Nppa*). Primers used are listed in **Table 1** (Gelinas *et al.*, 2008; Nawaito *et al.*, 2017). The abundance of mitochondrial calcium uptake 2 (*Micu2*), was performed by TaqMan assay (Assay ID: Mm00801666_g1, FAM-MGB) and normalized to *Gapdh* (Assay ID: Mm99999915_g1, VIC-MGB) as described previously (Nawaito *et al.*, 2019).

**Table 1.**
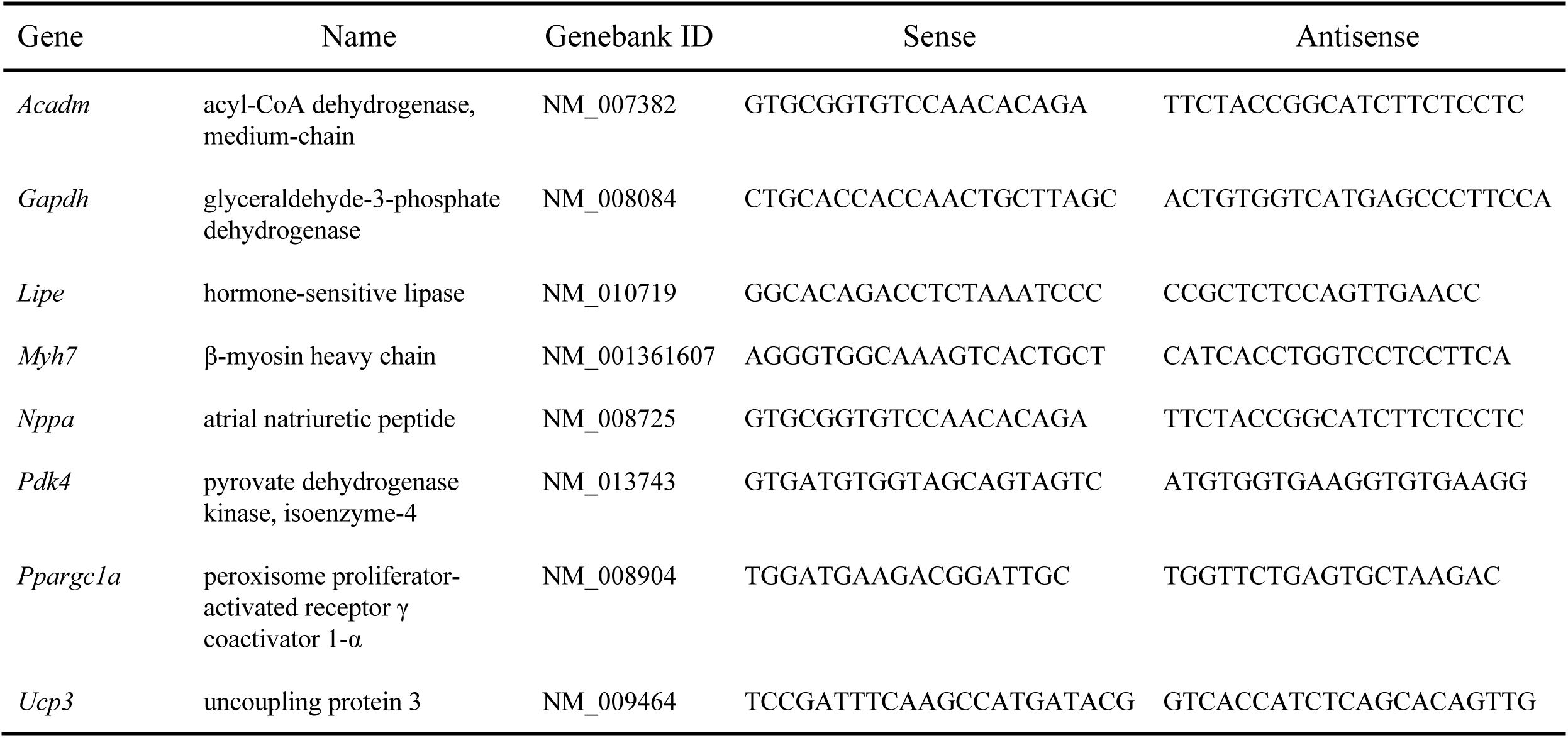
Primers used for quantitative PCR.

#### Biochemical analyses

Activities of the mitochondrial marker enzymes citrate synthase (CS), succinate dehydrogenase, isocitrate dehydrogenase (ICDH) and medium chain acyl-CoA dehydrogenase (MCAD) were measured spectrophotometrically in LV myocardium homogenates (Duda *et al.*, 2009; O’Shea *et al.*, 2009).

#### Immunoblotting

SDS-PAGE and immunoblotting were performed as described previously (Ruiz *et al.*, 2016) with a slight modification. Following electrophoretic transfer to 0.22 µm nitrocellulose, membranes were rinsed in PBS and fixed with glutaraldehyde (Connern & Halestrap, 1996) prior to blocking.

#### Statistical Analysis

Results are reported as the mean ± standard error (SEM). Statistical differences between two groups were tested using unpaired Student’s t-tests. For comparisons involving more than two groups, one-way ANOVA followed by a Bonferroni’s selected-comparison test was performed. For comparisons involving two independent variables, two-way ANOVA followed by Bonferroni’s post-test was performed. Survival curves were compared using Mantel-Cox log-rank tests. Results were considered statistically significant when *P*-values were less than 0.05. Statistical analyses were performed using GraphPad Prism version 8.2 for Mac OS X.

## Results

The development and general phenotypic characterization of MK2^−/−^ and MK2^+/−^ mice have been described elsewhere (Kotlyarov *et al.*, 1999) and the abundance of MK2 immunoreactivity in lysates prepared from the ventricular myocardium corelated with genotype (**Figure 1A**). At 12-weeks of age, MK2-deficient mice had similar body weight (MK2^+/+^: 29.7 ± 0.4 g, n = 141, MK2^−/−^: 28.7 ± 0.4 g, n = 143, *P* = 0.067) at 16 weeks of age they were characterized by a slight reduction in rectal temperature (MK2^+/+^: 33.1 ± 0.4 °C, n = 10, and MK2^−/−^: 32.0 ± 0.4 °C, n = 12, *P* < 0.05).

**Figure 1.**
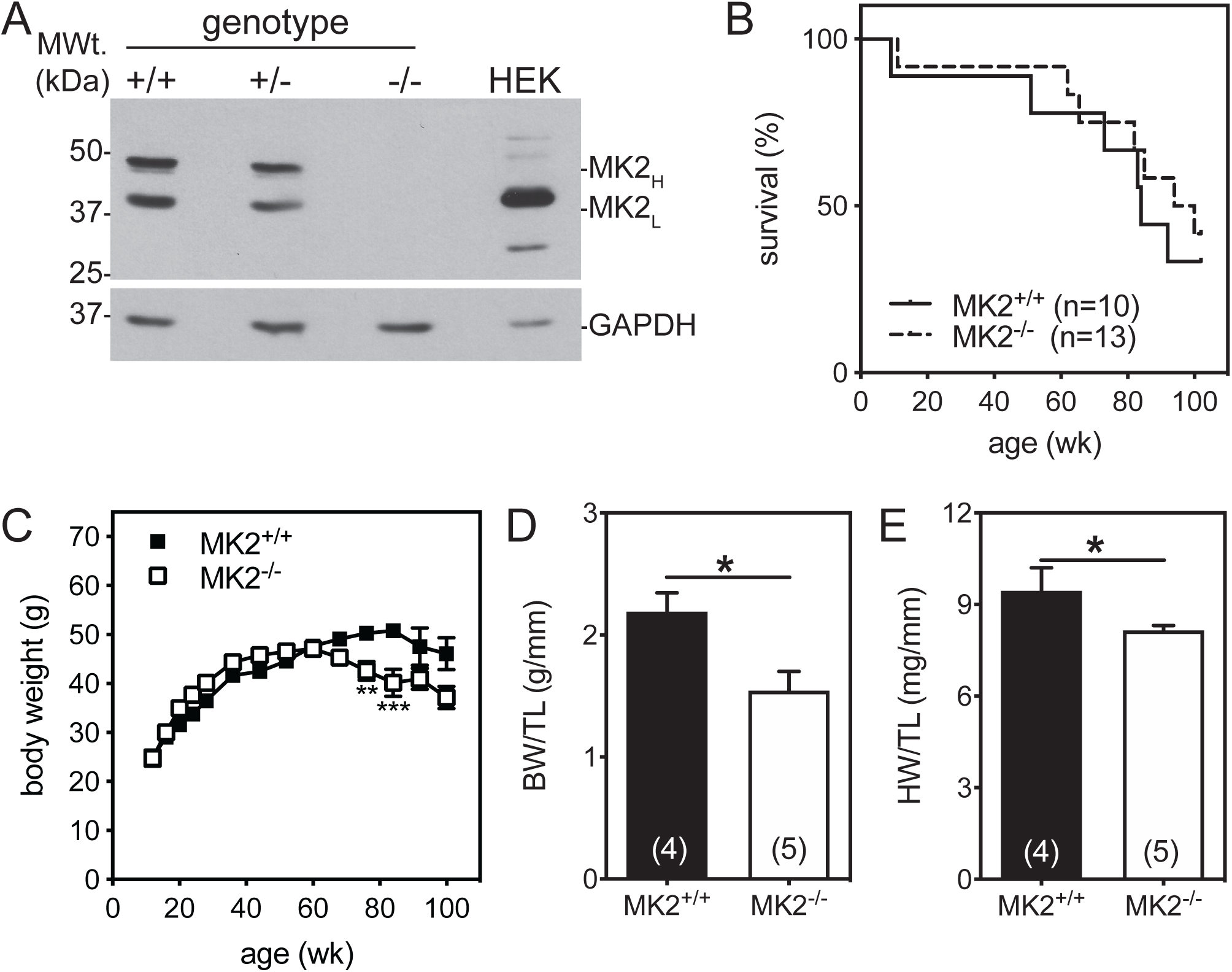
MK2-deficiency does not alter survival but enhances aging-dependent weight loss. **(A)** Representative image of heavy (H) and light (L) MK2 isoform immunoreactivity in heart lysates from MK2^+/+^, MK2^+/−^, and MK2^−/−^ mice. Numbers on left indicate molecular mass (in kDa) **(B)** Kaplan-Meier survival curves for MK2^+/+^ (n = 10) and MK2^−/−^ (n = 13) mice up to age 102 weeks. Mantel-Cox tests indicate the survival curves did not differ significantly (*P* = 0.580). **(C)** Changes in body weight in MK2^+/+^ and MK2^−/−^ mice up to 102 weeks of age. **(D)** body weight and **(E)** heart weight, both normalized to tibia length in 102-week-old MK2^+/+^ and MK2^−/−^ mice. Data are expressed as mean ± SEM. \**P* < 0.05, \*\**P* < 0.01, \*\*\**P* < 0.001.

### MK2^−/−^ mice show slight changes both in left and right ventricular function and bradycardia when assessed *in vivo*

#### Echocardiographic analyses

Echocardiographic imaging at 12-weeks of age revealed a prolonged R-R interval (MK2^+/+^: 176 ± 3 ms, n = 122, and MK2^−/−^: 199 ± 4 ms, n = 127, *P* < 0.0001) with modest differences in left ventricular (LV) structure and function in MK2-deficient mice **(Table 2)**. In MK2^−/−^ mice, several parameters reflecting LV structure were reduced significantly, including a 3% decrease in LV end-diastolic posterior wall thickness (LVPWd; MK2^+/+^: 0.736 ± 0.008 mm, n = 122, and MK2^−/−^: 0.714 ± 0.007 mm, n = 127, *P* < 0.05), a 4% decrease in LV end-diastolic diameter (LVDd; MK2^+/+^: 4.06 ± 0.03 mm, n = 122, and MK2^−/−^: 3.90 ± 0.03 mm, n = 127, *P* < 0.0001) and an 8% decrease in LV mass (MK2^+/+^: 111 ± 2 mg, n = 122, and MK2^−/−^: 102 ± 2 mg, n = 127, *P* < 0.001). LV ejection fraction was not significantly different in MK2^+/+^ (71.5 ± 0.7%, n = 122) and MK2^−/−^ (69.6 ± 0.9%, n = 127) mice. Consistent with a reduced LVDd and increased cardiac cycle length, cardiac output was reduced by 14% (MK2^+/+^: 11.4 ± 0.2 ml/min, n = 76, and MK2^−/−^: 9.78 ± 0.28 ml/min, n = 76, *P* < 0.0001) and both septal (S_S_; MK2^+/+^: 2.38 ± 0.04 cm/s, n = 122, and MK2^−/−^: 2.14 ± 0.04 cm/s, n = 127, *P* < 0.001) and lateral (S_L_; MK2^+/+^: 2.23 ± 0.04 cm/s, n = 122, and MK2^−/−^: 1.98 ± 0.04 cm/s, n = 127, *P* < 0.0001) wall systolic tissue velocities were decreased. Assessment of diastolic function revealed that the LV isovolumetric relaxation time, corrected for the differences in R-R interval, was prolonged by 16% in MK2^−/−^ mice (IVRT_c_; MK2^+/+^: 0.98 ± 0.03, n = 122, and MK2^−/−^: 1.14 ± 0.05, n = 127, *P* < 0.01). There were no differences in the flow velocity of either early (E wave; MK2^+/+^: 75.0 ± 1.0 cm/s, n = 122, and MK2^−/−^: 74.6 ± 1.4 cm/s, n = 127) or late (A wave; MK2^+/+^: 47.7 ± 0.7 cm/s, n = 109, and MK2^−/−^: 48.8 ± 0.7 cm/s, n = 120) left ventricular filling. In addition, E wave deceleration time was prolonged by 9.9% (EDT; MK2^+/+^: 35.2 ± 0.8 ms, n = 122, and MK2^−/−^: 38.7 ± 0.8 ms, n = 127, *P* < 0.01) and deceleration rate reduced by 9.6% (ED rate; MK2^+/+^: 22.8 ± 0.6 m/s^2^, n = 122, and MK2^−/−^: 20.6 ± 0.6 m/s^2^, n = 127, *P* < 0.01) in MK2^−/−^ hearts. Both septal and LV lateral mitral annular motion velocities (E_m_, A_m_) were reduced in MK2-deficient hearts, resulting in an increase in the ratio of transmitral early filling velocity to both lateral (lateral E/E_m_; MK2^+/+^: 36.7 ± 1.0, n = 110, and MK2^−/−^: 40.0 ± 1.1, n = 121, *P* < 0.05) and septal (septal E/E_m_; MK2^+/+^: 29.3 ± 0.6, n = 112, and MK2^−/−^: 32.7 ± 0.7, n = 120, *P* < 0.001) early diastolic mitral annular velocities, suggesting reduced LV compliance. Impaired LV compliance increases left atrial (LA) afterload and could result in LA enlargement. MK2^−/−^ mice actually showed a slight decrease in LA diameter during systole (LADs; MK2^+/+^: 2.24 ± 0.03 mm, n = 107, and MK2^−/−^: 2.12 ± 0.04 mm, n = 109, *P* < 0.05) with no changes in LA fractional shortening (LAFS; MK2^+/+^: 20.6 ± 0.5 %, n = 85, and MK2^−/−^: 20.4 ± 0.7%, n = 90). In addition, whereas a decrease in the ratio of the velocities of the systolic (S) and diastolic (D) waves of pulmonary venous flow would indicate impaired diastolic function, and this parameter is unaffected by increased R-R interval, the mean, upper, and lower pulmonary vein S/D ratios did not differ between MK2^+/+^ and MK2^−/−^ mice. In summary, MK2-deficiency did not alter LV systolic function but resulted in subtle signs of early diastolic dysfunction.

**Table 2.**
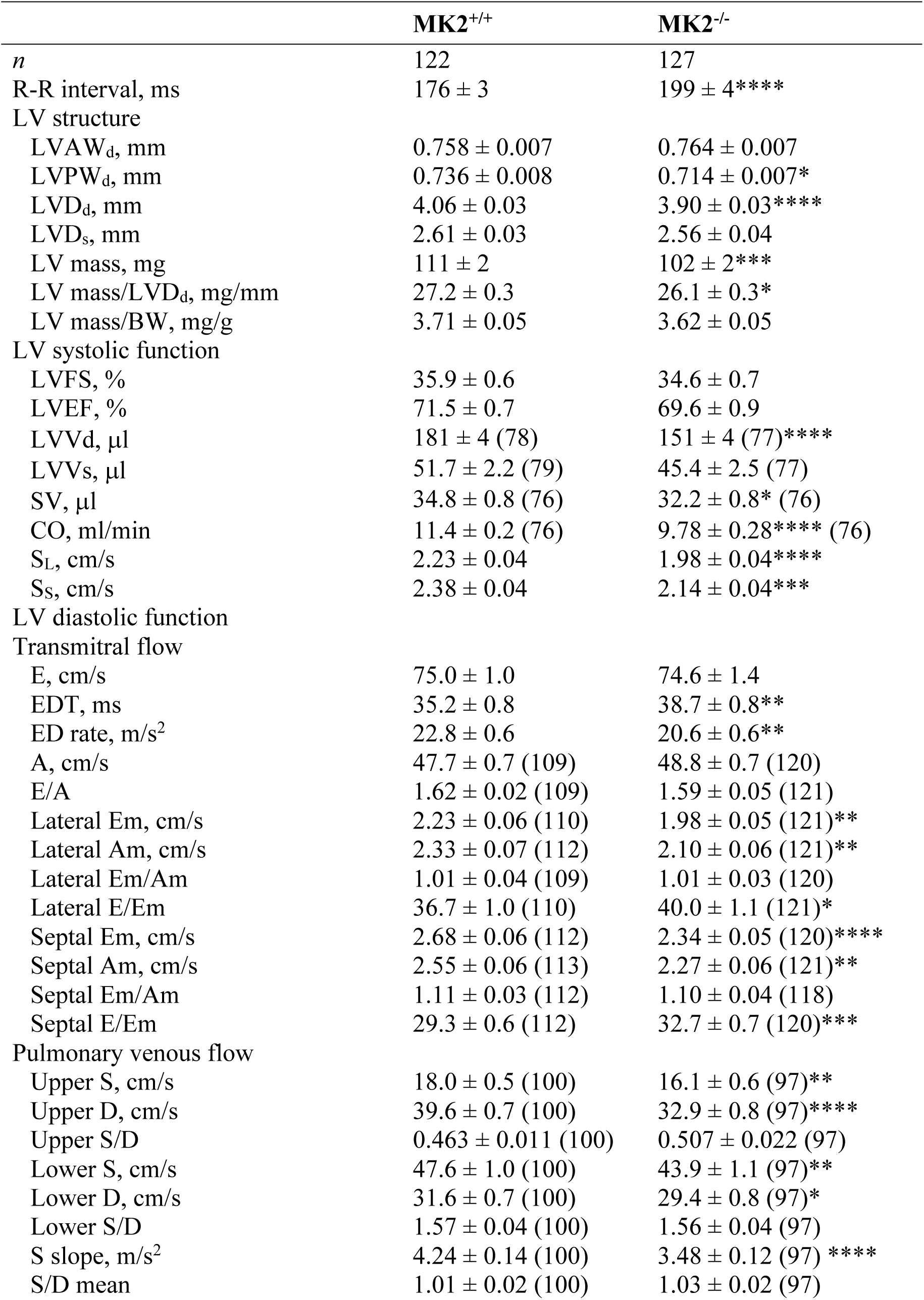
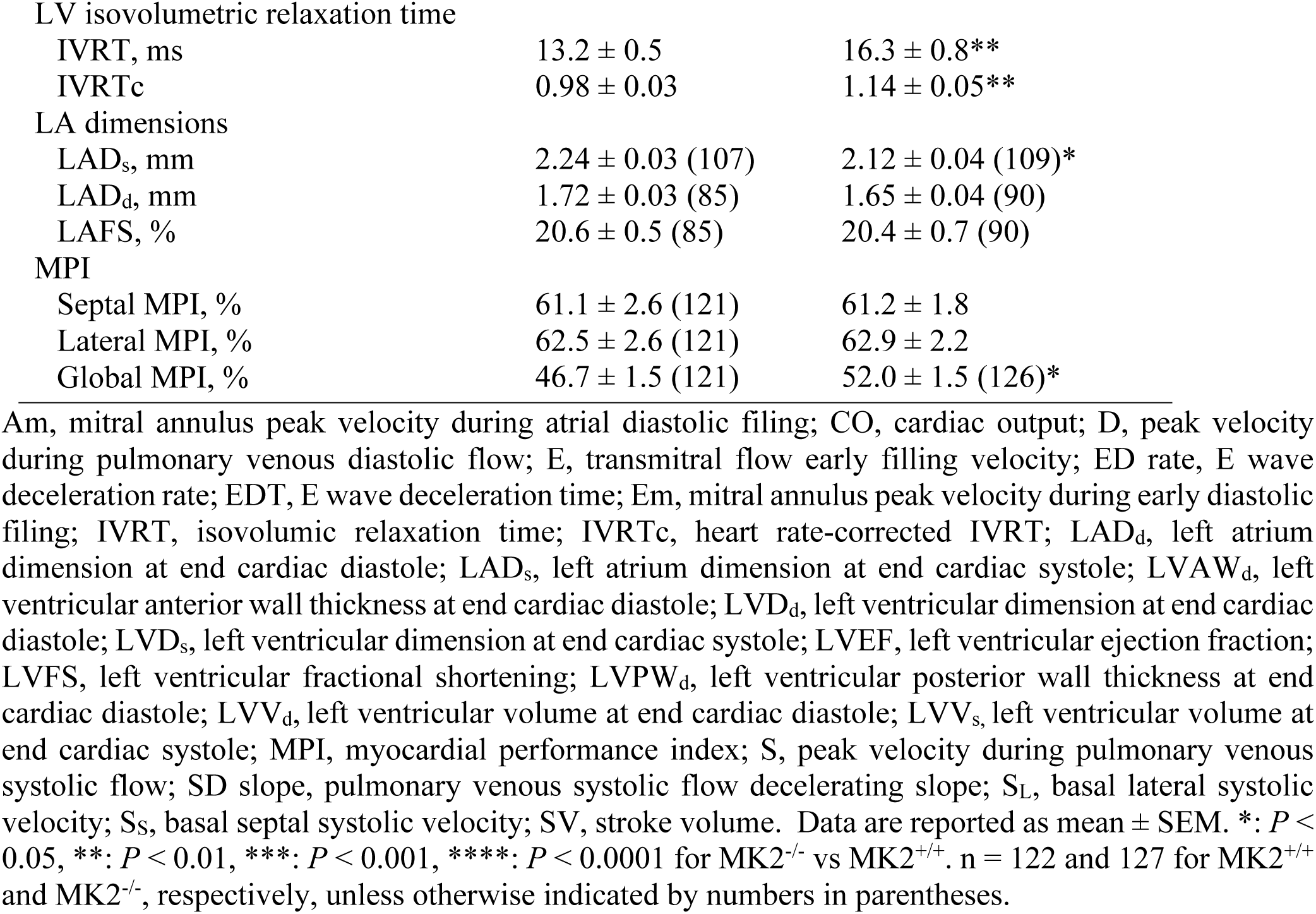
Echocardiographic parameters of LV structure and function in 12-week old MK2^+/+^ and MK2^−/−^ mice.

Right ventricular (RV) structure and function were also assessed by echocardiographic imaging **(Table 3)**. There were no major differences in terms of RV structure other than a small reduction in right ventricular end-diastolic diameter (RVDd; MK2^+/+^: 1.81 ± 0.02 mm, n = 78, and MK2^−/−^: 1.71 ± 0.03 mm, n = 77, *P* < 0.01). Tricuspid annular plane systolic excursion (TAPSE), which correlates closely with the right ventricular ejection fraction (Kaul *et al.*, 1984), was slightly reduced (TAPSE; MK2^+/+^: 1.26 ± 0.02 mm, n = 78, and MK2^−/−^: 1.21 ± 0.01 mm, n = 77, *P* < 0.01) in MK2^−/−^ mice as was the RV systolic tricuspid annular velocity (S_R_; MK2^+/+^: 3.32 ± 0.07 cm/s, n = 78, and MK2^−/−^: 2.81 ± 0.07 cm/s, n = 77, *P* < 0.0001). In terms of RV diastolic function, peak velocities of early (E_T_; MK2^+/+^: 30.3 ± 0.8 cm/s, n = 77, and MK2^−/−^: 30.7 ± 1.2 cm/s, n = 77) and late (A_T_; MK2^+/+^: 42.4 ± 0.9 cm/s, n = 68, and MK2^−/−^: 42.6 ± 1.3 cm/s, n = 66) RV filling, in addition to the E_T_ wave deceleration time and deceleration rate (E_T_D rate), were similar in MK2^+/+^ and MK2^−/−^ hearts. Both early (E_m_; MK2^+/+^: 3.34 ± 0.10 cm/s, n = 71, and MK2^−/−^: 2.72 ± 0.08 cm/s, n = 65, *P* < 0.0001) and late (A_m_; MK2^+/+^: 3.22 ± 0.10 cm/s, n = 71, and MK2^−/−^: 2.83 ± 0.11 cm/s, n = 72, *P* < 0.05) diastolic lateral tricuspid annular motion velocities were reduced in MK2-deficient hearts, resulting in a 16% increase in the ratio of trans-tricuspid early filling velocity to early diastolic tricuspid annular velocity (E/E_m_; MK2^+/+^: 9.09 ± 0.30, n = 70, and MK2^−/−^: 10.5 ± 0.4, n = 65, *P* < 0.01). Hence, echocardiographic assessment revealed that the deletion of MK2 resulted in changes in RV function similar to those observed in the LV.

**Table 3.** Echocardiographic parameters of RV structure and function in 12-week old MK2^+/+^ and MK2^−/−^ mice.

|  | MK2<sup>+/+</sup> | MK2<sup>-/-</sup> |
| --- | --- | --- |
| *n* | 78 | 77 |
| RV structure |  |  |
| RVAW<sub>d</sub>, mm | 0.316 ± 0.006 | 0.299 ± 0.007 |
| RVD<sub>d</sub>, mm | 1.81 ± 0.02 | 1.71 ± 0.03** |
| RVAW<sub>d</sub>/RVD<sub>d</sub> | 0.175 ± 0.003 | 0.175 ± 0.004 |
| TAPSE, mm | 1.26 ± 0.02 | 1.21 ± 0.01* |
| RV systolic function |  |  |
| S<sub>R</sub>, cm/s | 3.32 ± 0.07 | 2.81 ± 0.07**** |
| RV diastolic function |  |  |
| Transtricuspid flow |  |  |
| E<sub>t</sub>, cm/s | 30.3 ± 0.8 (77) | 30.7 ± 1.2 |
| E<sub>t</sub>DT, ms | 29.3 ± 1.3 | 29.4 ± 1.3 |
| E<sub>t</sub>D rate, m/s<sup>2</sup> | 10.2 ± 0.4 (77) | 10.2 ± 0.5 |
| A<sub>t</sub>, cm/s | 42.4 ± 0.9 (68) | 42.6 ± 1.3 (66) |
| E<sub>t</sub>/A<sub>t</sub> | 0.693 ± 0.024 | 0.727 ± 0.045 |
| Lateral Em, cm/s | 3.34 ± 0.10 (71) | 2.72 ± 0.08 (65)**** |
| Lateral Am, cm/s | 3.22 ± 0.10 (71) | 2.83 ± 0.11 (65)* |
| Lateral Em/Am | 1.09 ± 0.04 (71) | 1.08 ± 0.07 (65) |
| Lateral E/Em | 9.09 ± 0.30 (70) | 10.5 ± 0.4 (65)** |
| Pulmonary artery flow |  |  |
| AT, ms | 21.9 ± 0.7 (75) | 23.1 ± 0.8 (76) |
| AT/RVET | 26.7 ± 0.7 (75) | 24.9 ± 0.7 (76) |
| RV MPI |  |  |
| TVCO, ms | 102 ± 2 (77) | 115 ± 2 (76)**** |
| RVET, ms | 82.2 ± 1.4 (75) | 92.6 ± 1.7**** |
| Lateral MPI, % | 39.2 ± 0.8 | 40.4 ± 0.9 |
| Global MPI, % | 24.5 ± 1.2 (74) | 27.6 ± 1.5 (73) |

Am, tricuspid annulus peak velocity during late (atrial) diastolic filing; At, transtricuspid flow late (atrial) filling velocity; AT, pulmonary arterial flow acceleration time; Em, tricuspid annulus peak velocity during early diastolic filing; Et, transtricuspid flow early filling velocity; EtD rate, transtricuspid early filling deceleration rate; EtDT, transtricuspid early filling deceleration time; MPI, myocardial performance index; RVAW<sub>d</sub>, right ventricular anterior wall thickness at end cardiac diastole; RVD<sub>d</sub>, right ventricular diameter at end cardiac diastole; RVET, right ventricular ejection time; S<sub>R</sub>, right ventricular lateral wall systolic velocity; TAPSE, tricuspid anulus plane systolic excursion; TVCO; tricuspid valve closure to opening time. Data are reported as mean ± SEM. \*: *P* < 0.05, \*\*: *P* < 0.01, \*\*\*: *P* < 0.001, \*\*\*\*: *P* < 0.0001 for MK2<sup>-/-</sup> vs MK2<sup>+/+</sup>. *n* = 78 and 77 for MK2<sup>+/+</sup> and MK2<sup>-/-</sup>, respectively, unless otherwise indicated by numbers in parentheses.

We then examined the possible effect(s) of an MK2-deficiency on cardiac aging. Mantel-Cox tests indicated that the survival curves for wild-type (10) and MK2-deficient (13) male littermates did not differ significantly when assessed up to 102 weeks of age (**Figure 1B**, *P* = 0.580). Although both MK2^+/+^ and MK2^−/−^ mice initially appeared to age similarly, body weight in MK2^−/−^ mice peaked at 60 weeks-of-age and then began to decrease whereas that of wild type mice peaked at around 80 weeks-of-age (**Figure 1C**). At 102 weeks of age, the body weight of surviving MK2^−/−^ mice was 29% lower than that of surviving MK2^+/+^ mice (MK2^+/+^: 2.29 ± 0.16 g/mm, n = 4, and MK2^−/−^: 1.55 ± 0.16 g/mm, n = 5, *P* < 0.05) when normalized to tibia length (**Figure 1D**). At 102 weeks of age, surviving MK2^−/−^ mice also showed a 14% decrease (MK2^+/+^: 9.46 ± 0.75 mg/mm, n = 4, and MK2^−/−^: 8.15 ± 0.16 mg/mm, n = 5, *P* < 0.05) in heart weight normalized to tibia length (**Figure 1E**). Food and water consumption of 88-week-old MK2-deficient mice did not differ significantly from that of wild-type littermates (Data Not Shown). Cardiac structure and function were assessed in MK2^+/+^ and MK2^−/−^ mice by echocardiography every 4 (weeks 12 – 28) or 8 (weeks 28 - 100) weeks until 102 weeks of age (**Table 4**). MK2^−/−^ mice displayed a prolonged R-R interval throughout the study; however, in the second year of life the difference no longer reached significance. Throughout the duration of the study, left ventricular ejection fraction and fractional shortening in MK2-deficient mice did not differ significantly from that of wild type litter mates indicating systolic function was unaffected. When examined at 3 months of age, MK2-deficient mice showed signs of reduced compliance, suggesting early diastolic dysfunction (**Table 2**); however, this condition did not progress with age (**Table 4**). Specifically, peak E and A wave velocities, as well as regional E/E_m_ ratios, did not differ significantly. In these smaller groups of mice, E wave deceleration times did not differ significantly whereas the deceleration rate tended to be significantly slower. Furthermore, in MK2-deficient mice, aging was not associated with left atrial dilation or changes in the pulmonary venous flow S/D ratio, which are hallmarks of diastolic dysfunction. However, although the left atria were significantly smaller in MK2^−/−^ mice, this difference was lost with age, suggesting there may have been some degree of atrial remodeling in these mice over time. In summary, the absence of MK2 was associated with weight loss but no age-related progressive deterioration in systolic or diastolic function.

**Table 4.** MK2-deficiency does not result in a progressive decline in cardiac function.

|  | < 6 mo |  | 6 – 12 mo |  | 12 – 18 mo |  | 18 – 24 mo |  |
| --- | --- | --- | --- | --- | --- | --- | --- | --- |
|  | MK2<sup>+/+</sup> | MK2<sup>-/-</sup> | MK2<sup>+/+</sup> | MK2<sup>-/-</sup> | MK2<sup>+/+</sup> | MK2<sup>-/-</sup> | MK2<sup>+/+</sup> | MK2<sup>-/-</sup> |
| *n* | 10 | 13 | 10 | 12 | 9 | 12 | 6 | 9 |
| R-R interval, ms | 156±6 | 184±8* | 148±3 | 178±5** | 166±5 | 187±4 | 158±13 | 176±9 |
| LV structure |  |  |  |  |  |  |  |  |
| LVAW<sub>d</sub>, mm | 0.716±0.026 | 0.727±0.016 | 0.834±0.017 | 0.850±0.017 | 1.070±0.039 | 0.943±0.031** | 1.100±0.035 | 1.003±0.033 |
| LVPW<sub>d</sub>, mm | 0.718±0.026 | 0.724±0.018 | 0.803±0.016 | 0.800±0.021 | 0.873±0.040 | 0.869±0.080 | 0.923±0.034 | 0.902±0.023 |
| LVD<sub>d</sub>, mm | 3.98±0.05 | 3.84±0.12 | 4.20±0.04 | 4.27±0.06 | 4.34±0.09 | 4.56±0.06 | 4.32±0.10 | 4.33±0.12 |
| LV mass, mg | 102±6 | 98.2±6.4 | 132±3 | 139±5 | 175±7 | 175±11 | 184±6 | 171±8 |
| LV mass/LVD<sub>d</sub>, mg/mm | 25.6±1.2 | 25.2±1.0 | 31.4±0.7 | 32.3±0.9 | 40.6±1.8 | 38.4±2.4 | 42.7±1.7 | 39.4±0.9 |
| LV systolic function |  |  |  |  |  |  |  |  |
| LVFS, % | 38.1±1.2 | 35.3±1.8 | 39.9±0.5 | 36.2±1.2 | 36.6±1.3 | 33.5±2.2 | 37.4±1.0 | 34.6±1.2 |
| LVEF, % | 74.3±1.4 | 69.5±2.3 | 76.4±0.5 | 71.5±1.6 | 72.3±1.7 | 67.6±2.7 | 73.9±1.3 | 70.1±1.7 |
| SL, cm/s | 2.58±0.11 | 2.33±0.10 | 2.86±0.08 | 2.44±0.09** | 1.76±0.09 | 1.71±0.06 | 1.82±0.10 | 1.67±0.12 |
| SS, cm/s | 2.28±0.11 | 2.08±0.08 | 2.74±0.10 | 2.36±0.08* | 1.97±0.08 | 1.75±0.07 | 1.97±0.08 | 1.72±0.11 |
| LV diastolic function |  |  |  |  |  |  |  |  |
| Transmitral flow |  |  |  |  |  |  |  |  |
| E, cm/s | 60.4±1.9 | 53.2±3.9 | 68.1±1.4 | 62.4±1.5 | 73.7±3.3 | 70.4±2.9 | 79.2±5.7 | 73.2±2.6 |
| EDT, ms | 34.3±1.2 | 40.4±2.2 | 34.0±1.5 | 40.7±1.4 | 39.7±1.3 | 42.5±1.9 | 33.4±3.7 | 38.8±2.8 |
| ED rate, m/s<sup>2</sup> | 19.7±1.2 | 14.3±1.3** | 20.9±0.7 | 16.0±0.8* | 19.2±1.3 | 17.1±0.7 | 25.6±2.9 | 20.1±12* |
| A, cm/s | 44.3±2.1 | 49.0±2.4 | 47.4±1.9 | 51.9±1.8 | 42.1±4.0 | 50.0±4.0 | 39.2±4.0 | 52.1±4.2 |
| E/A | 1.31±0.04 | 1.06±0.07 | 1.40±0.06 | 1.20±0.05 | 1.97±0.30 | 1.68±0.32 | 2.23±0.29 | 1.44±0.16 |
| Lateral Em, cm/s | 2.43±0.12 | 1.75±0.12** | 2.52±0.14 | 1.98±0.15* | 1.77±0.12 | 1.87±0.19 | 1.67±0.13 | 1.78±0.20 |
| Lateral Em/Am | 0.796±0.065 | 0.723±0.050 | 0.818±0.044 | 0.664±0.061 | 0.938±0.057 | 0.97±0.11 | 0.813±0.034 | 0.99±0.13 |
| Lateral E/Em | 26.3±1.7 | 25.7±1.8 | 28.6±1.8 | 36.1±2.0 | 46.2±4.5 | 41.7±4.9 | 46.8±1.3 | 43.5±5.2 |
| Septal Em, cm/s | 2.30±0.24 | 1.95±0.15 | 2.53±0.17 | 2.11±0.19 | 2.44±0.19 | 2.1±0.13 | 2.22±0.30 | 2.27±0.26 |
| Septal Em/Am | 0.858±0.097 | 0.819±0.108 | 0.944±0.083 | 0.782±0.087 | 1.34±0.14 | 1.27±0.12 | 1.16±0.12 | 1.16±0.07 |
| Septal E/Em | 30.2±2.9 | 28.1±2.2 | 29.0±2.1 | 34.1±2.6 | 33.4±2.4 | 36.4±2.9 | 36.8±3.0 | 34.7±4.7 |
| Pulmonary venous flow |  |  |  |  |  |  |  |  |
| S/D mean | 1.28±0.10 | 1.22±0.09 | 1.19±0.09 | 1.40±0.06 | 1.33±0.13 | 1.65±0.16 | 1.22±0.14 | 1.60±0.12 |
| LV isovolumetric relaxation time |  |  |  |  |  |  |  |  |
| IVRTc | 1.34±0.05 | 1.36±0.07 | 1.22±0.06 | 1.43±0.06 | 1.11±0.06 | 1.36±0.05* | 0.94±0.05 | 1.19±0.11 |
| LA dimensions |  |  |  |  |  |  |  |  |
| LAD<sub>s</sub>, mm | 2.06±0.10 | 1.67±0.06** | 2.37±0.06 | 2.40±0.06 | 2.85±0.15 | 3.00±0.08 | 3.08±0.11 | 3.16±0.10 |
| LAD<sub>d</sub>, mm | 1.68±0.10 | 1.31±0.07* | 1.92±0.05 | 1.94±0.06 | 2.33±0.16 | 2.45±0.09 | 2.64±0.12 | 2.63±0.12 |
| LAFS, % | 18.9±1.1 | 22.6±1.4 | 19.3±0.8 | 19.2±1.1 | 19.0±1.5 | 18.9±1.3 | 14.2±1.2 | 17.3±1.7 |
| MPI |  |  |  |  |  |  |  |  |
| Septal MPI, % | 72.1±5.0 | 67.7±3.4 | 69.5±4.7 | 71.1±2.6 | 44.0±4.1 | 48.3±3.2 | 60.9±7.4 | 48.4±4.4 |
| Lateral MPI, % | 69.9±5.3 | 64.0±2.5 | 69.5±5.1 | 73.6±4.4 | 42.2±4.1 | 57.4±4.4 | 59.3±7.0 | 51.9±4.3 |
| Global MPI, % | 41.0±3.3 | 57.2±2.9** | 33.0±3.2 | 49.2±3.9** | 25.2±2.8 | 42.6±2.6** | 37.9±6.6 | 47.3±4.7 |

#### Heart rate and electrocardiography in conscious mice

As mentioned above, echocardiographic imaging, obtained under isoflurane anesthesia, revealed that the R-R interval was prolonged in MK2^−/−^ mice **(Table 2)**, indicating a lower heart rate in these mice (HR; MK2^+/+^: 341 ± 20 beats/min, n = 122, and MK2^−/−^: 306 ± 17 beats/min, n = 127, *P* < 0.0001). Analysis by surface electrocardiogram revealed that when mice under isoflurane anesthesia were challenged with a single dose of isoproterenol (0.1 mg/kg) the heart rate of MK2^−/−^ mice and MK2^+/+^ mice increased to similar levels, whereas HR declined more rapidly in MK2^−/−^ mice (**Figure 2A**). However, isoflurane lowers HR (Constantinides & Murphy, 2016) and could, possibly, affect MK2^+/+^ and MK2^−/−^ mice differently. Hence, we next evaluated cardiac electrical activity in conscious unrestrained mice by radio telemetry over a period of 48 hours. Overall, the mean R-R interval in MK2^−/−^ mice was 7% longer in comparison to their littermate MK2^+/+^ mice even in the absence of anesthetic (MK2^+/+^: 115 ± 3 ms, n = 6, and MK2^−/−^: 123 ± 2 ms, n = 6, *P* < 0.05) **(Table 5)**. Stated otherwise, the heart rate in MK2^+/+^ mice was 538 ± 11 bpm (n = 6) whereas that of MK2^−/−^ mice was 496 ± 8 bpm (n = 6; *P* < 0.05) **(Table 5, Figure 2C)**. In addition, the duration of the P-R segment was longer in MK2^−/−^ mice (MK2^+/+^: 22.6 ± 0.4 ms, n = 5, and MK2^−/−^: 25.9 ± 1.0 ms, n = 4, *P* < 0.05) whereas the P wave duration, QRS duration, and QTc were unaffected **(Table 5)**. An examination of the circadian rhythm pattern of heart rate revealed MK2-deficient mice showed prolonged R-R intervals **(Figure 2D,E)** and P-R segments **(Figure 2F)** during both light and dark cycles. We next performed a non-linear analysis of heart rate variability by plotting the R-R interval in the form of Poincaré plots (R-R_n_ versus R-R_n+1_), fitting the data to an ellipse, and determining the standard deviation on the short (SD1) and long (SD2) axes of the ellipse. SD1 reflects short term variability in heart rate whereas SD2 reflects both short term and long term variability (Shaffer & Ginsberg, 2017). The SD1/SD2 ratio was significantly larger in MK2-deficient mice **(Figure 2G)**. Both prolonged P-R segment, reflecting atrioventricular nodal conduction times, and increased short term variability in heart rate suggest autonomic regulation of heart rate is altered in MK2-deficient mice.

**Figure 2.**
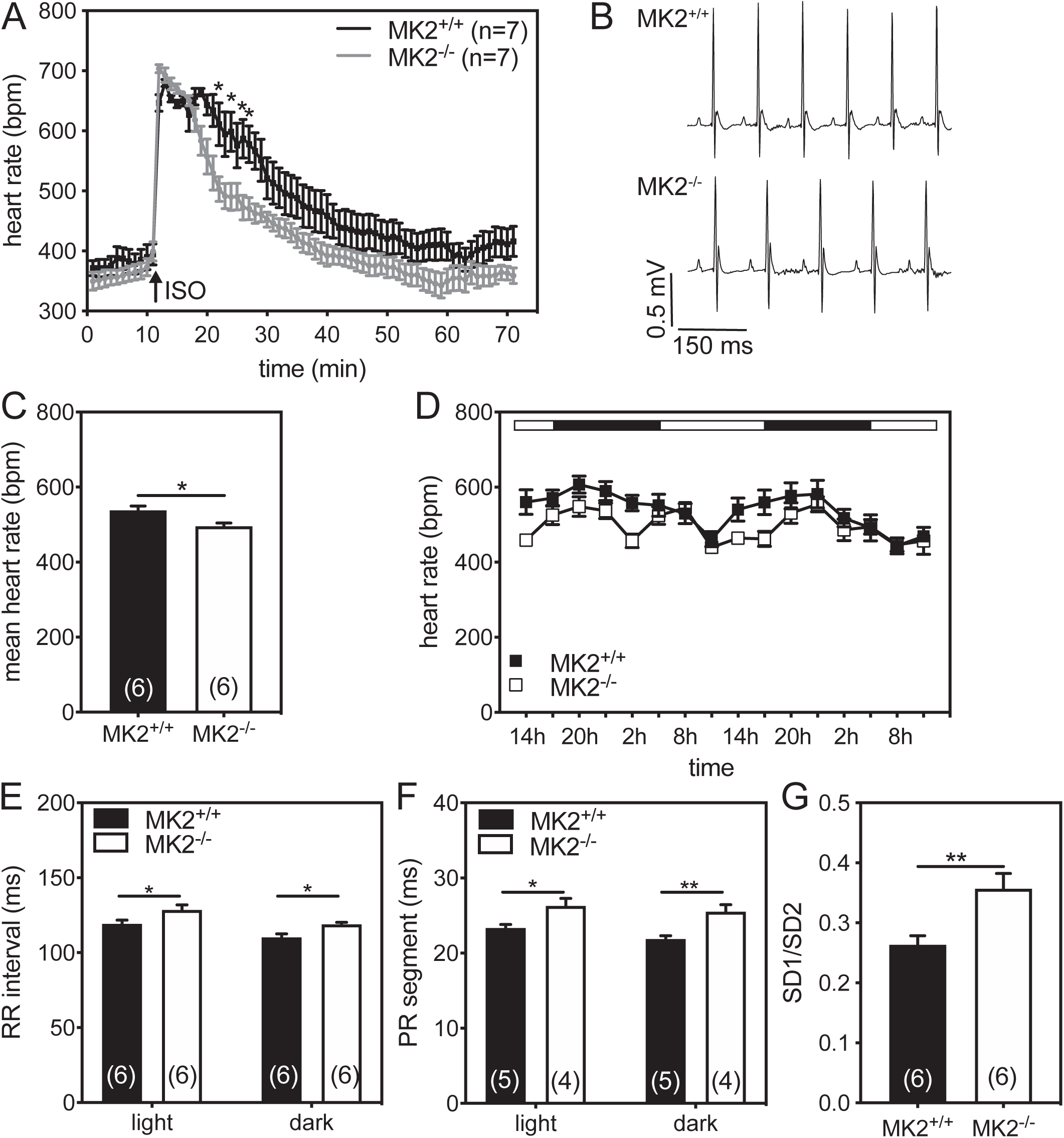
MK2^−/−^ mice exhibited a slower heart rate *in vivo*. **(A)** *In vivo* heart rate (HR) assessment by surface electrocardiography (ECG) in MK2^+/+^ (n = 7) and MK2^−/−^ (n = 7) mice anesthetized with isoflurane. Arrow indicates injection of isoproterenol (ISO, 0.1 mg/kg). \**P* < MK2^+/+^ vs. MK2^−/−^, repeated-measures two-way ANOVA with Bonferroni post-test. **(B)** Representative electrocardiograms obtained by radio telemetry in conscious unrestrained MK2^+/+^ (upper) and MK2^−/−^ (lower) mice. **(C)** Mean heart rates obtained from ECGs recorded over a 48-h time period by radio telemetry in conscious unrestrained MK2^+/+^ (n = 6) and MK2^−/−^ (n = 6) mice. Data was extracted and analyzed in 3 h intervals and then averaged separately for each mouse prior to calculating the mean heart rate for each group. **(D)** Heart rates obtained from ECGs recorded over a 48-h time period by radio telemetry in conscious unrestrained MK2^+/+^ (n = 6) and MK2^−/−^ (n = 6) mice. Data was extracted and analyzed in 3 h intervals. Mice were maintained on a 12:12 hour light:dark cycle (6h00:18h00) with the bar at the top of the graph indicating light and dark cycles. The circadian rhythm is clearly visible. **(E)** R-R interval and **(F)** P-R segment duration obtained from ECGs recorded over a 48-h time period by radio telemetry in conscious unrestrained MK2^+/+^ (n = 6) and MK2^−/−^ (n = 6) mice. Data was extracted and analyzed in 3 h intervals and then values for light and dark cycles averaged separately for each mouse prior to calculating the mean heart rate for each group. **(G)** Mean 48 h SD1/SD2 ratios for MK2^+/+^ (n = 6) and MK2^−/−^ (n = 6) mice where SD1 represents the standard deviation on the short axis and SD2 represents the standard deviation on the long axis of an ellipse fit to a plot of R-R_n_ versus R-R_n+1_. Data are expressed as mean ± SEM. \**P* < 0.05, \*\**P* < 0.01.

**Table 5.** ECG parameters obtained from conscious 12-week old MK2^+/+^ and MK2^−/−^ mice.

|  | **MK2<sup>+/+</sup>** | **MK2<sup>-/-</sup>** |
| --- | --- | --- |
| *n* | 6 | 6 |
| R-R interval, ms | 115 ± 3 | 123 ± 2* |
| Heart rate, bpm | 538 ± 11 | 496 ± 8* |
| P-R interval, ms | 31.3 ± 0.5 (5) | 35.2 ± 0.9 (4)** |
| P wave duration, ms | 8.70 ± 0.22 (5) | 9.25 ± 0.42 (4) |
| P-R segment, ms | 22.6 ± 0.4 (5) | 25.9 ± 1.0* (4) |
| QRS duration, ms | 8.89 ± 0.48 | 8.67 ± 0.69 |
| P wave amplitude, mV | 0.0132 ± 0.0027 (5) | 0.0127 ± 0.0022 (4) |
| QT interval, ms | 46.1 ± 0.6 | 49.1 ± 1.1* |
| QTc | 43.4 ± 0.4 | 44.1 ± 1.1 |

Data are reported as mean ± SEM. \*: $P < 0.05$ , \*\*: $P < 0.01$ . *n* = 6 unless otherwise indicated by numbers in parentheses.

### Calcium transients are similar in isolated ventricular cardiomyocytes from MK2^+/+^ and MK2^−/−^ mice

In light of the differences in R-R interval and diastolic function observed in MK2^−/−^ mice (**Table 2, Table 4, Figure 2**), we examined the effect of an MK2-deficiency on the amplitude and decay of Ca^2+^ transients evoked by field stimulation (2 Hz, **Figure 3A to C**) or superfusion with caffeine (10 s, 10 μM, **Figure 3D to G**) in isolated adult cardiac ventricular myocytes. Representative transients from MK2^+/+^ and MK2^−/−^ myocytes are respectively shown in **Figure 3, Panels A and D**. The mean data shows in both conditions that neither the amplitude of the cytosolic Ca^2+^ transient nor the time to 90% transient decay for Ca^2+^-transients differed significantly in cardiomyocytes from MK2^+/+^ and MK2^−/−^ mice after field stimulation (**Figure 3B,C**) or following caffeine superfusion (**Figure 3E,F**). In addition, the fractional release of Ca^2+^, calculated by dividing the amplitude of the depolarization-induced Ca^2+^ transient by that of the caffeine-induced transient, was also similar in MK2^+/+^ and MK2^−/−^ myocytes (**Figure 3G**). Hence, the absence of MK2 did not significantly alter the Ca^2+^ content of cardiomyocyte SR or its capacity for Ca^2+^ release and reuptake.

**Figure 3.**
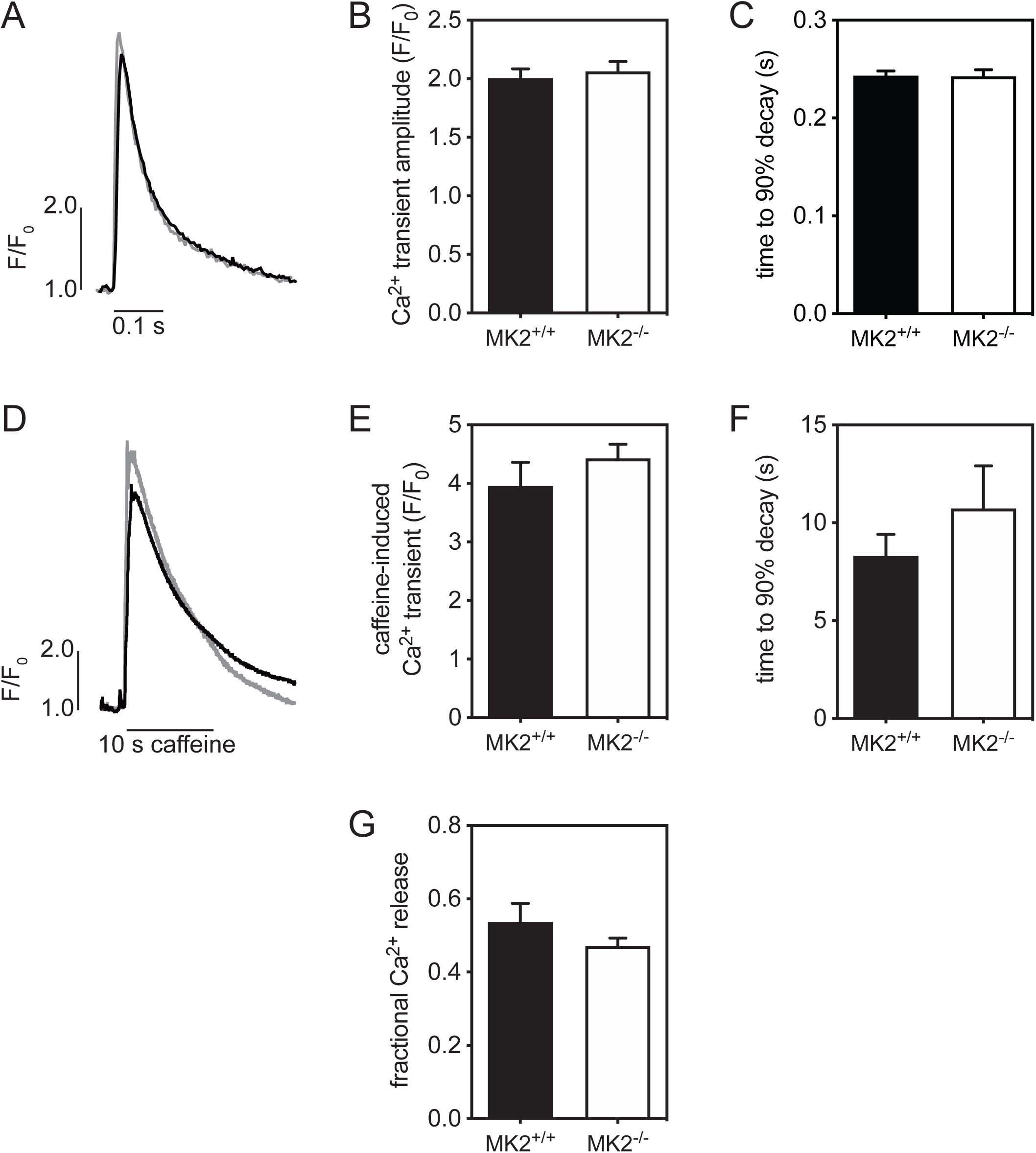
MK2 deficiency does not alter calcium transients or SR calcium content in adult cardiac ventricular myocytes. Representative pacing-induced **(A)** and caffeine-induced **(D)** Ca^2+^ transients in ventricular myocytes isolated from twelve-week-old MK2^+/+^ (black) and MK2^−/−^ (grey) mice, respectively. Mean Ca^2+^ transient amplitudes evoked by pacing **(B)** and caffeine **(E)** in ventricular myocytes MK2^+/+^ and MK2^−/−^ ventricular myocytes. Panels **C** and **F** show time to 90% decay for pacing- or caffeine-induced transients, respectively. (**G)** Mean values for fractional SR calcium release (pacing-induced transient divided by caffeine-induced transient) in MK2^+/+^ and MK2^−/−^ ventricular myocytes. Data shown are mean ± SEM from a total of eight myocytes from four separate cell preparations.

### Heart rate normalizes when hearts from MK2^−/−^ mice are perfused *ex vivo* in working mode

To determine if the observed bradycardia involved mechanisms intrinsic to the MK2-deficient heart, we next examined heart function in isolated hearts. The heart simultaneously uses multiple substrates for energy production (Khairallah *et al.*, 2004); hence, cardiac function was examined in the presence of 1) exogenous carbohydrates and 2) exogenous carbohydrates plus palmitate as energy sources. First, in contrast to observations of HR made *in vivo*, when hearts were perfused *ex vivo* in a semi-recirculating working mode, heart rates in MK2^+/+^ and MK2^−/−^ mice were similar (MK2^+/+^: 463 ± 27 beats/min, n = 8, and MK2^−/−^: 491 ± 19 beats/min, n = 10; **Table 6, Figure 4**). Our assessment of heart function *ex vivo* included evaluating the response to isoproterenol wherein hearts from MK2^+/+^ and MK2^−/−^ mice were perfused in working mode for 20 min, after which isoproterenol (ISO, 10 nM) was added to the perfusion buffer for an additional 20 min. Hearts from both MK2^−/−^ and MK2^+/+^ mice responded to the presence of ISO in the perfusate with an increase in HR and this increase was of similar magnitude (MK2^+/+^: 590 ± 18 beats/min, n = 7, and MK2^−/−^: 575 ± 12 beats/min, n = 10; **Table 6, Figure 4**). When perfused with buffer containing ISO, values for most LV function parameters, especially related to systolic function and cardiac flow, remained unchanged in both MK2^+/+^ and MK2^−/−^ hearts, although LV developed pressure increased significantly in MK2-deficient, but not wild type hearts. Similarly, ISO induced a significant increase in the maximum rate of relaxation (-d*P*/d*t*) in MK2-deficient, but not wild type hearts, compared to baseline. The mouse heart normally derives more than half of its energy requirements from fatty acid oxidation. No functional differences were observed between MK2^+/+^ and MK2^−/−^ hearts when perfused in the presence of palmitate (0.6 mM), although it is worth noting that with the addition of palmitate to the perfusing buffer, there were more functional changes in MK2-deficient hearts, relative to carbohydrates alone as the exogenous energy source, than observed in wild type hearts (**Table 6**). Overall, it was interesting to note that, whereas MK2-deficient hearts were bradycardic when assessed *in vivo*, when cardiac function was assessed *ex vivo* in an isolated working heart preparation, no difference in heart rate was observed.

**Figure 4.**
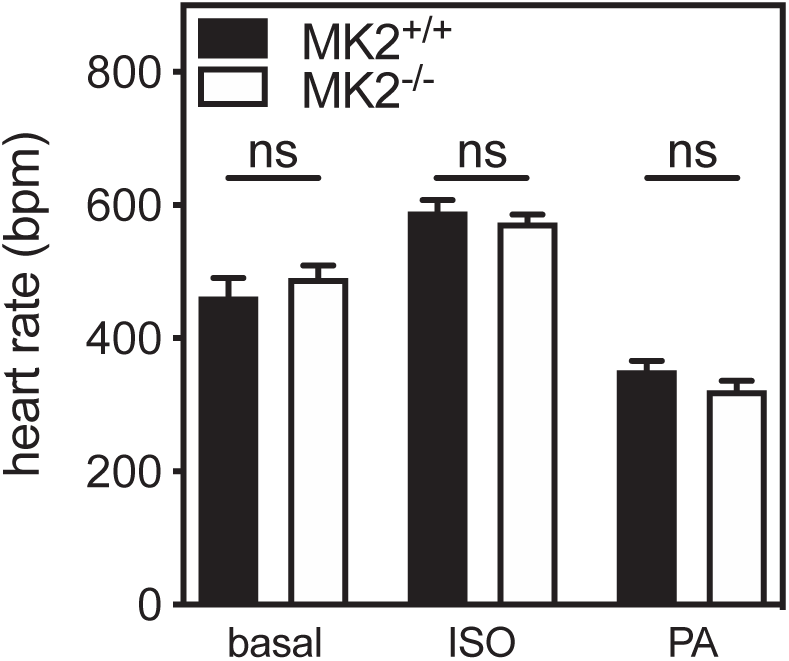
Heart rates do not significantly differ when MK2^+/+^ and MK2^−/−^ hearts are perfused *ex vivo*. Hearts rate in isolated working hearts from MK2^+/+^ and MK2^−/−^ mice perfused with modified Krebs-Henseleit buffer alone **(basal)**, buffer containing 10 nM isoproterenol **(ISO**), or mM palmitic acid **(PA)**. Data shown are mean ± SEM of 7 – 11 heart preparations. The actual sample size for each group is as described in Table 6. ns, not significantly different by two-way ANOVA with Bonferroni post-tests.

**Table 6.**
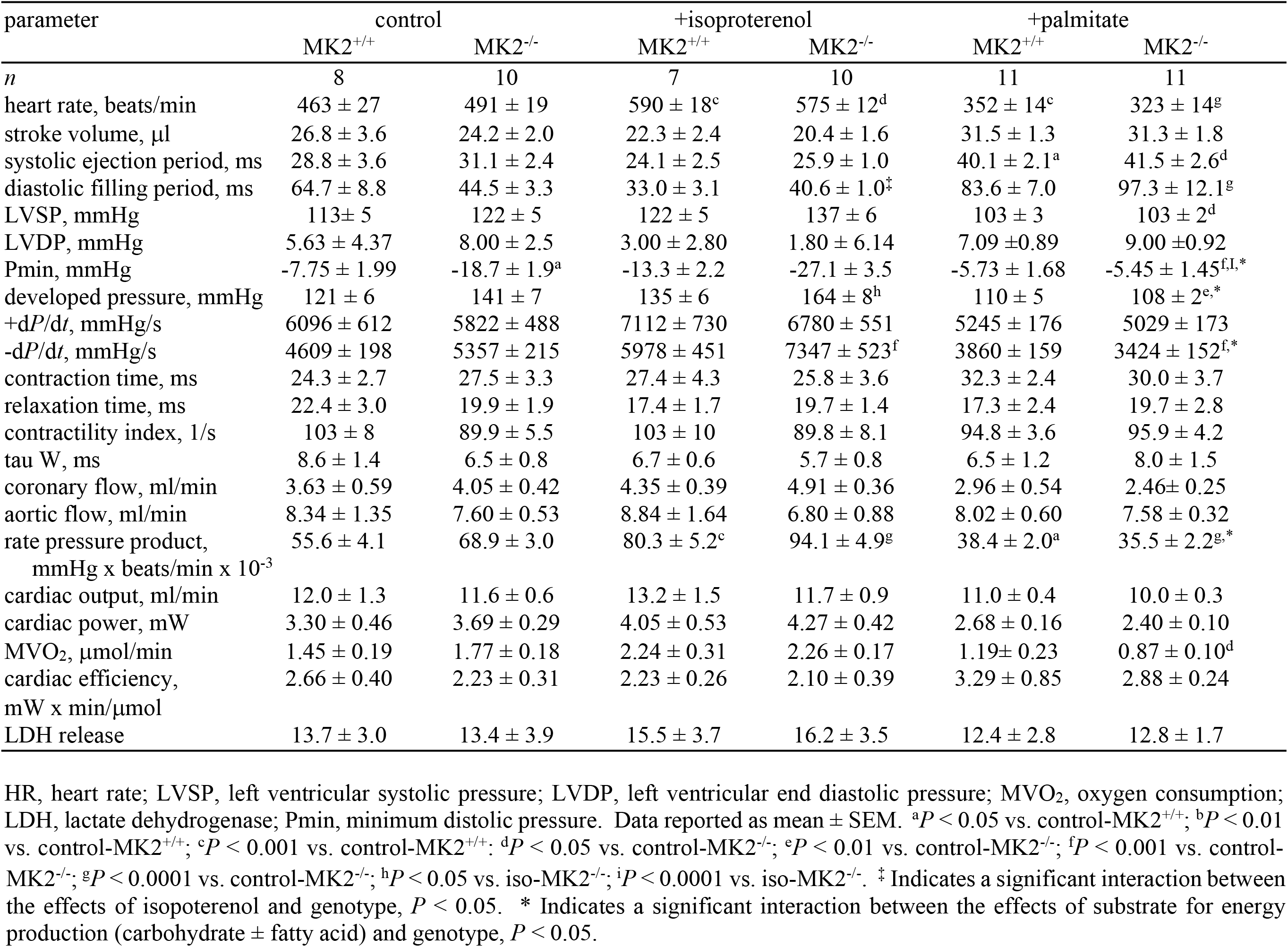
Functional parameters of isolated working MK2^+/+^ and MK2^−/−^ mouse hearts.

### MK2^−/−^-mice are characterized by changes in transcript abundance for metabolic genes and increased sensitivity to PTP opening

Although it is estimated that mitochondria contribute marginally to Ca^2+^ removal during global Ca^2+^ transients in healthy cardiomyocytes, mitochondria occupy ≈ 30% of the volume of a cardiomyocyte and may therefore influence diastolic Ca^2+^. In this regard, and due to the age-dependent weight loss observed in MK2-deficient mice, we examined certain aspects of mitochondria function. First, we evaluated function in isolated mitochondria by measuring oxygen consumption in state 3 and state 4. Compared with mitochondria from wild-type hearts, those from MK2^−/−^ hearts showed no difference in mitochondrial respiration, irrespective of substrates used, or in mitochondrial generation of reactive oxygen species, specifically hydrogen peroxide (**Table 7**). However, mitochondria from MK2^−/−^ hearts showed a decreased sensitivity to MPTP opening as evidenced by the greater cumulative calcium load needed to trigger MPTP opening (**Figure 5A**). Interestingly, this occurred despite no significant changes in the abundance of immunoreactivity of different proteins putatively involved in the cardiac MPTP complex, namely cyclophilin-D (CypD), voltage-dependent anion channel (VDAC), and adenine nucleotide translocase (ANT: **Figure 5C**). Cytochrome c oxidase subunits I (COX-I) and 4 (COX-IV) served as loading controls for mitochondrial protein **(Figure 5C)**. As single nucleotide polymorphisms in the mitochondrial calcium uptake 2 (*MICU2/EFHA1*) gene have been associated with changes in P-R segment duration (Verweij *et al.*, 2014), transcript levels for *Micu2* were assessed by qPCR and found to be reduced in hearts from MK2^−/−^ mice **(Figure 5B)**.

**Figure 5.**
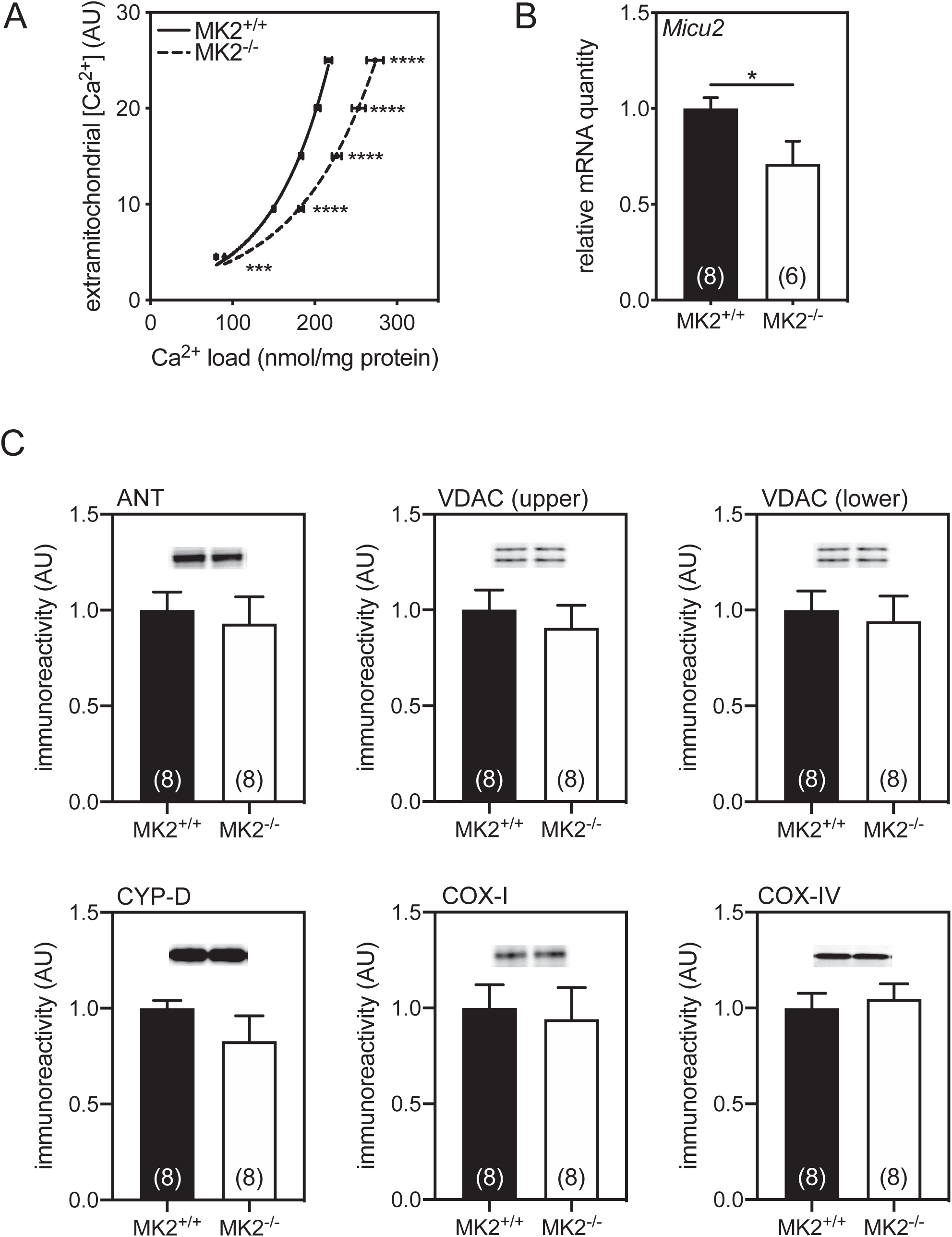
Calcium-sensitivity in isolated mitochondria from 12-week-old MK2^−/−^ showed delayed mPTP opening compared to their littermate counterparts. Subsarcolemmal mitochondria were isolated from MK2^−/−^ and MK2^+/+^ hearts and assessed for their ability to buffer extramitochondrial calcium. **(A)** Low calcium loads are sustainable by mitochondria; however, when a threshold calcium load is attained, a large and abrupt increase in extramitochondrial calcium signifies opening of the mitochondrial permeability transition pore (MPTP). **(B)** The abundance of the mitochondrial Ca^2+^ uptake 2 (*Micu2*) mRNA, a regulatory subunit of the mitochondrial inner membrane Ca^2+^ uniporter, in the ventricular myocardium. **(C)** The abundance of immunoreactivity of the different proteins responsible for mitochondrial permeability transition. Cyp-D, cyclophilin-D; VDAC, voltage-dependent anion channel; ANT, adenine nucleotide translocase. Cytochrome c oxidase subunits 1 (COX-I) and 4 (COX-IV) immunoreactivity served as loading controls. AU, arbitrary units. Data are expressed as mean ± SEM (n = 8). * *P* < 0.05, *** *P* < 0.001, **** *P* < 0.0001, MK2^+/+^ *vs.* MK2^−/−^.

**Table 7.** Mitochondrial parameters from MK2^−/−^ and MK2^+/+^ hearts.

|  |  | MK2<sup>+/+</sup> | MK2<sup>-/-</sup> |
| --- | --- | --- | --- |
| *n* |  | 8 | 8 |
| Mitochondrial Respiration, atoms O/mg/min |  |  |  |
| Glutamate + Malate |  |  |  |
|  | State 3 | 130 ± 6 | 141 ± 12 |
|  | State 4 | 33.0 ± 2.0 | 38.6 ± 2.4 |
|  | RCR | 4.3 ± 0.4 | 3.6 ± 0.2 |
| Pyruvate + Malate |  |  |  |
|  | State 3 | 412 ± 17 | 455 ± 24 |
|  | State 4 | 55.2 ± 4.7 | 62.0 ± 3.2 |
|  | RCR | 8.2 ± 0.6 | 7.4 ± 0.5 |
| Palmitoyl-Carnitine + Malate |  |  |  |
|  | State 3 | 441 ± 21 | 399 ± 38 |
|  | State 4 | 49.7 ± 5.9 | 45.0 ± 4.6 |
|  | RCR | 8.8 ± 1.3 | 10.1 ± 1.0 |
| Succinate + Rotenone |  |  |  |
|  | State 3 | 467 ± 29 | 423 ± 26 |
|  | State 4 | 167 ± 18 | 166 ± 15 |
|  | RCR | 3.7 ± 0.7 | 2.7 ± 0.2 |
| Mitochondrial enzymatic activities |  |  |  |
| CS activity, μmol/mg/min |  | 1.76 ± 0.12 | 1.66 ± 0.06 |
| MCAD activity, μmol/mg/min |  | 0.18 ± 0.01 | 0.18 ± 0.01 |
| SD activity, μmol/mg/min |  | 0.015 ± 0.001 | 0.02 ± 0.001* |
| ICDH activity, mmol/mg/min |  | 1.29 ± 0.15 | 1.08 ± 0.07 |
| Mitochondrial H<sub>2</sub>O<sub>2</sub> production rate, nmol/min/mg |  |  |  |
|  | State 3 | 0.07 ± 0.01 | 0.07 ± 0.01 |
|  | State 4 | 0.37 ± 0.03 | 0.38 ± 0.02 |

Data are mean ± SEM. RCR, respiratory control ratio; CS, citrate synthase; MCAD, medium chain acyl-CoA dehydrogenase, SD, succinate dehydrogenase; ICDH, isocitrate dehydrogenase. \* p<0.05, MK2<sup>+/+</sup> vs. MK2<sup>-/-</sup>.

At the metabolic level, although serum glucose, free fatty acids, triglycerides, and ketone bodies levels did not differ between MK2^−/−^ mice and MK2^+/+^ mice (Data Not Shown), the abundance of mRNAs encoding medium-chain acyl-CoA dehydrogenase (MCAD), hormone-sensitive lipase (HSL), peroxisome proliferator-activated receptor γ coactivator 1α (PGC1α), and uncoupling protein 3 (UCP3) were significantly higher in ventricular myocardium from MK2^−/−^ mice compared to controls (**Figure 6**). In addition, the activity of mitochondrial enzymes isocitrate dehydrogenase, citrate synthase, and MCAD was similar between both groups, whereas that of succinate dehydrogenase activity was increased by 30% in mitochondria from MK2^−/−^ hearts (SD; MK2^+/+^: 0.015 ± 0.001 μmol/mg/min, n = 8, and MK2^−/−^: 0.020 ± 0.001 μmol/mg/min, n = 8, *P* < 0.05, **Table 7**).

**Figure 6.**
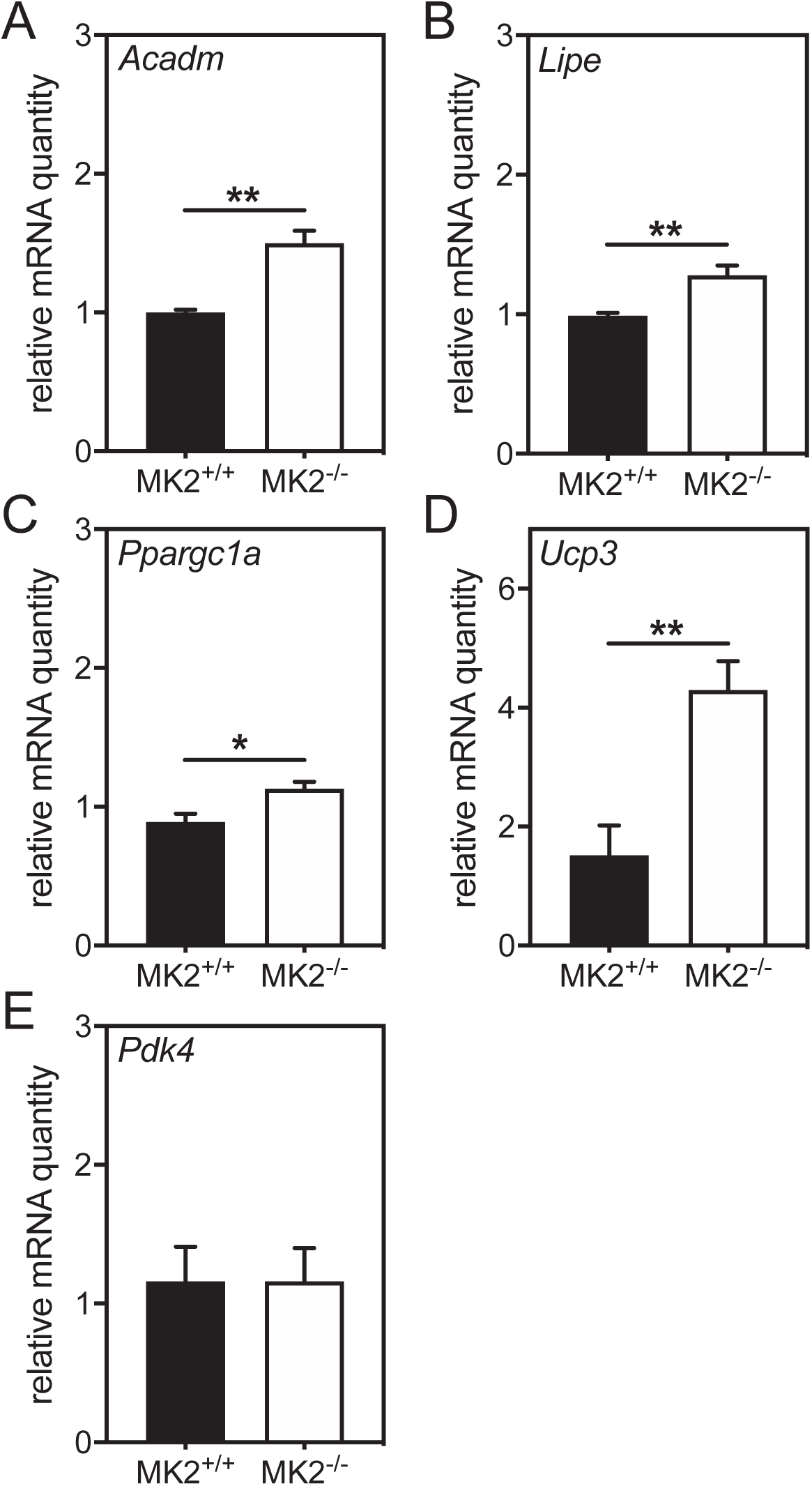
MK2^−/−^ mice showed changes in specific mitochondrial associated markers. The abundance of mRNA for genes encoding **(A)** medium-chain acyl-CoA dehydrogenase (*Acadm*), **(B)** hormone-sensitive lipase (*Lipe*), **(C)** peroxisome proliferator-activated receptor γ coactivator 1α (*Ppargc1a*), **(D)** uncoupling protein 3 (*Ucp3*), and **(E)** pyruvate dehydrogenase kinase 4 (*Pdk4*) was evaluated on RNA isolated from the ventricular myocardium. Data were normalized to the abundance of *Gapdh* mRNA. Values shown were normalized to that of a representative MK2^+/+^ mouse and expressed as mean ± SE. n = 4 mice/group. \**P* < 0.05, ** *P* < 0.01, MK2^+/+^ *vs.* MK2^−/−^. * *P* < 0.05, ** *P* < 0.01, MK2^+/+^ *vs.* MK2^−/−^.

### Pressure overload-induced myocardial remodeling is delayed, but not prevented in MK2-deficient mice

Given 1) the above-mentioned marginal effects of MK2 deletion on baseline cardiac function, 2) the cardioprotective effects of an MK2-deficiency in a streptozotocin model of diabetes (Ruiz *et al.*, 2016), and 3) increased afterload induced by constriction of the transverse aorta (TAC) both activates p38 MAPKs and increases phosphorylation of hsp25 at residues phosphorylated by MKs 2, 3, and 5 (serine-15, serine-82) (Esposito *et al.*, 2001; Dingar *et al.*, 2010), we next assessed the effect of MK2-deficiency on cardiac remodeling in response to TAC. Twelve-week-old male MK2-deficient and wild-type littermate male mice underwent TAC and were sacrificed 2 or 8 weeks of post-surgery, as described previously (Nawaito *et al.*, 2017). Invasive hemodynamic assessment revealed that TAC induced increases in both peak systolic arterial pressure and peak left ventricular pressure that did not differ significantly between TAC-MK2^+/+^ and TAC-MK2^−/−^ groups **(Table 8, Figure 7A)** with an average overall increase in peak systolic arterial pressure of 40 mmHg. In addition, echocardiographic imaging of the aortic arch revealed comparable flow velocities at the site of constriction in TAC-MK2^+/+^ and TAC-MK2^−/−^ groups **(Table 9)**. Hence, an increase in afterload was similarly established in both MK2^+/+^ and MK2^−/−^ mice. Two weeks post-TAC, echocardiographic imaging revealed left ventricular mass increased by 47% (sham: 103 ± 3 mg, n = 17; TAC: 151 ± 8 mg, n = 16, *P* < 0.0001) in MK2^+/+^ mice, which was attenuated to an increase of 9% in MK2-deficient mice that did not reach statistical significance (sham: 112 ± 6 mg, n = 14; TAC: 128 ± 7 mg, n = 15, *P* > 0.05). Similarly, heart weight to tibia length ratios (HW/TL) were increased by 37% in MK2^+/+^ (sham: 67.9 ± 3.9 mg/cm, n = 8; TAC: 92.8 ± 5.1 mg/cm, n = 11, *P* < 0.001) and by only 7.8% in MK2^−/−^ mice (sham: 66.9 ± 3.8 mg/cm, n= 10; TAC: 77.1 ± 2.1 mg/cm, n = 10, *P* > 0.05). The increase in HW/TL ratio observed in MK2-deficient mice was significantly less than that observed in wild type mice (*P* < 0.05, **Figure 7B**). Pathological left ventricular hypertrophy is associated with molecular remodeling that includes the re-expression of cardiac fetal genes atrial natriuretic peptides (*Nppa*) and β-myosin heavy chain (*Myh7*) (Akashi *et al.*, 2007). *Nppa* and *Myh7* mRNA levels were quantified by qPCR two weeks post-TAC. Although hypertrophy was attenuated two weeks post-TAC in MK2 deficient mice, *Nppa* **(Figure 8A)** and *Myh7* **(Figure 8B)** mRNA levels in MK2^+/+^ and MK2^−/−^ hearts increased to similar levels. Eight weeks post-TAC, LV mass was similarly increased in MK2^+/+^ mice by 64% (sham: 106 ± 3 mg, n = 11; TAC: 174 ± 14 mg, n = 11, *P* < 0.0001) and in MK2^−/−^ mice by 50% (sham: 109 ± 3 mg, n = 11; TAC: 164 ± 8 mg, n = 10, *P* < 0.001) **(Table 9)**. Hence, although hypertrophy was attenuated in MK2-deficient mice two weeks post-TAC, this represented a delay in the hypertrophic response.

**Figure 7.**
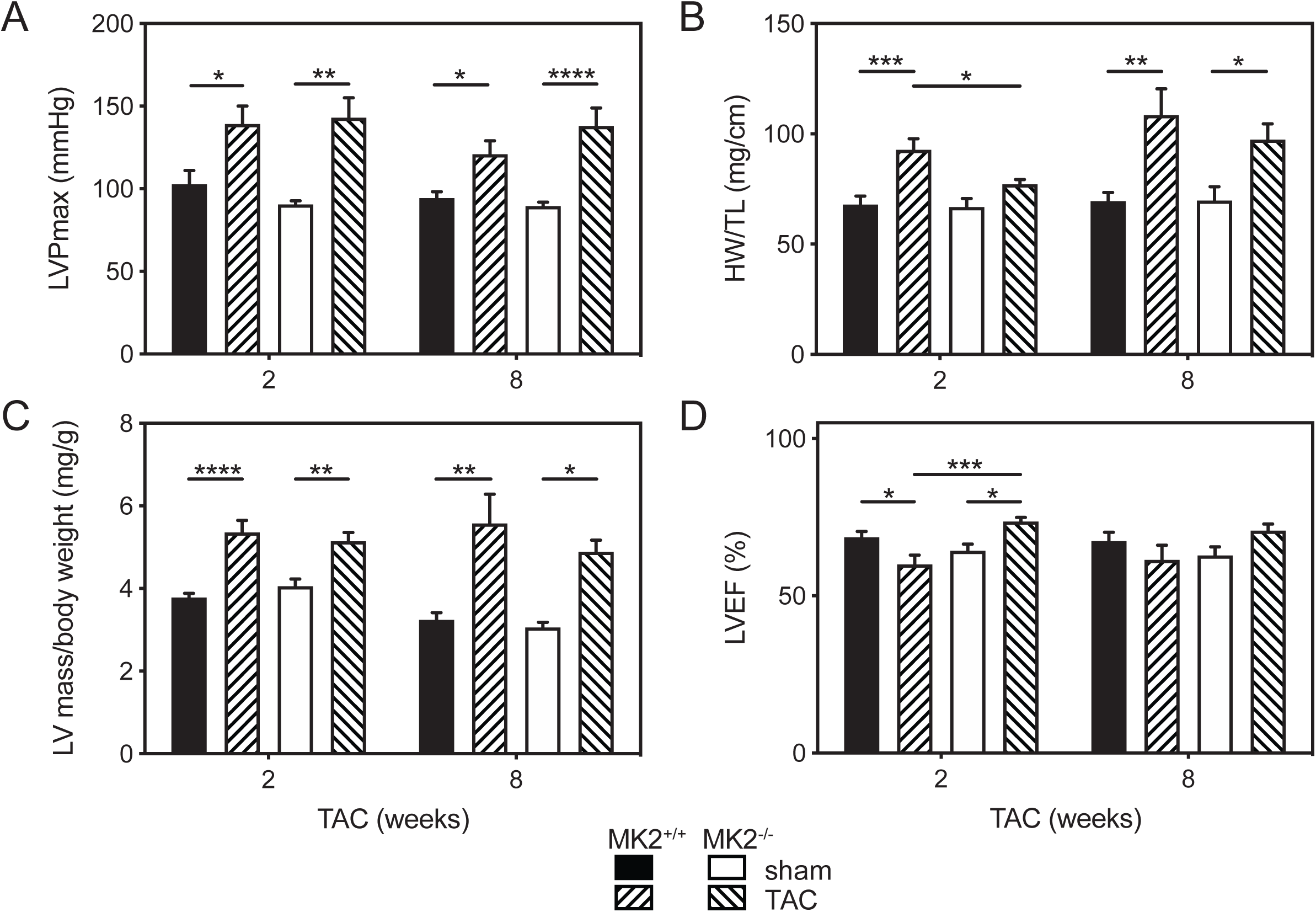
Hypertrophy is delayed in MK2^−/−^ hearts subjected to transverse aortic constriction (TAC). Pressure overload was induced in MK2^−/−^ and wild-type littermate control (MK^+/+^) mice by TAC, and mice were euthanized 2 and 8 wk post-surgery. Sham-operated (sham) animals underwent the identical surgical procedure; however, the aorta was not constricted. Left ventricular (LV) maximum developed pressure (LVPmax; **A**), heart weight-to-tibia length ratio (HW/TL; **B**), LV mass-to-body weight ratio (LV mass/body weight; **C**), and left ventricular ejection fraction (LVEF, **D**). Heart weight refers to the mass of the whole heart (minus atria) as determined gravimetrically. LV mass was determined by echocardiography. Data are expressed as means ± SE; n = 6 – 17 mice. *, *P* < 0.05, \*\**P* < 0.01, \*\*\**P* < 0.001, and \*\*\*\**P* < 0.001 by two-way ANOVA with Bonferroni’s multiple-comparison post-test.

**Figure 8.**
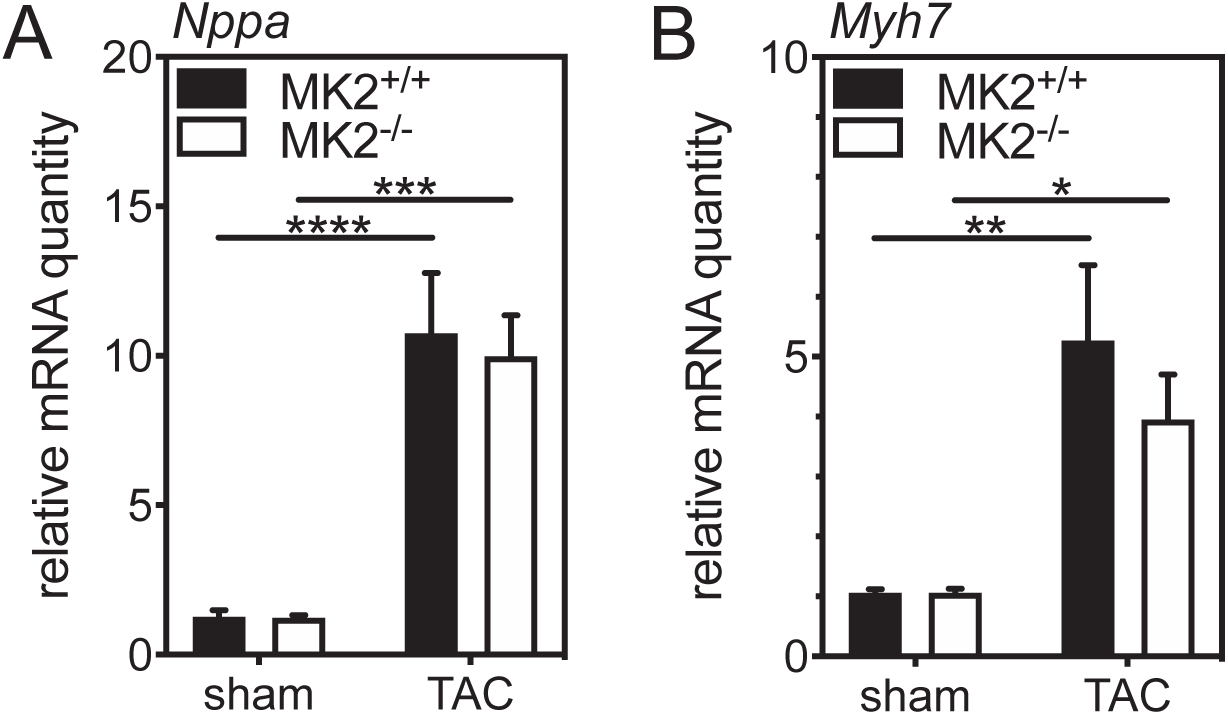
MK2-deficiency does not alter the TAC-induced increase in fetal gene expression. Pressure overload was induced in MK2^−/−^ and wild-type littermate control (MK2^+/+^) mice by TAC, and mice were euthanized 2 wk post-surgery. Sham animals underwent the identical surgical procedure; however, the aorta was not constricted. Total RNA was isolated and the abundance of atrial natriuretic peptide (*Nppa*) and **(B)** β-myosin heavy chain (*Myh7*) mRNA was quantified by quantitative PCR and normalized to *Gapdh* mRNA levels. Data are expressed as means ± SE; n = 5-6. \**P* < 0.05, **P 0.01, \*\*\**P* < 0.001, and \*\*\*\**P* < 0.001 by two-way ANOVA with Bonferroni’s multiple-comparison post-test.

**Table 8.**
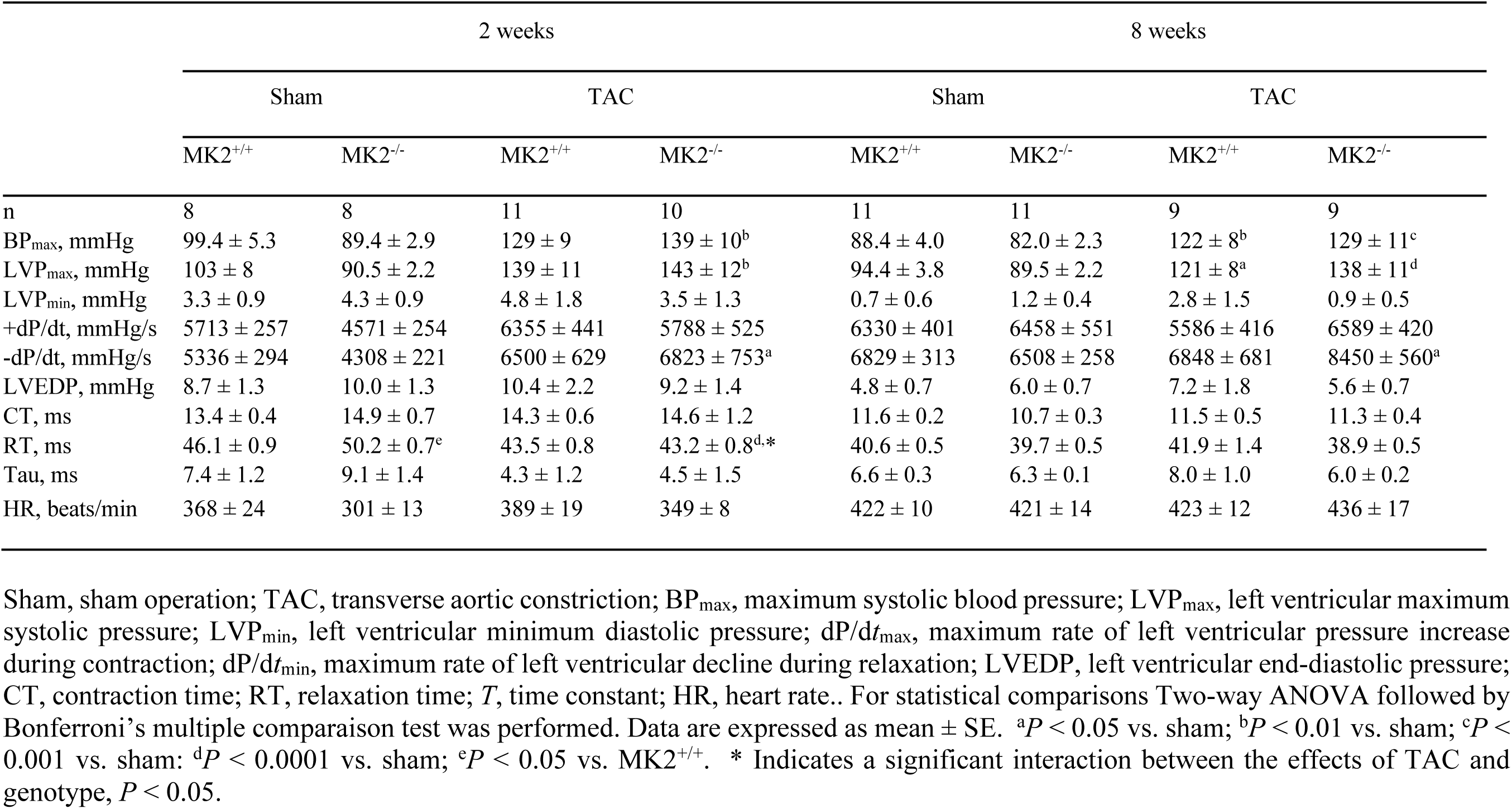
Hemodynamic parameters for MK2+/+ and MK2−/− mice 2- and 8-weeks post-TAC.

**Table 9.**
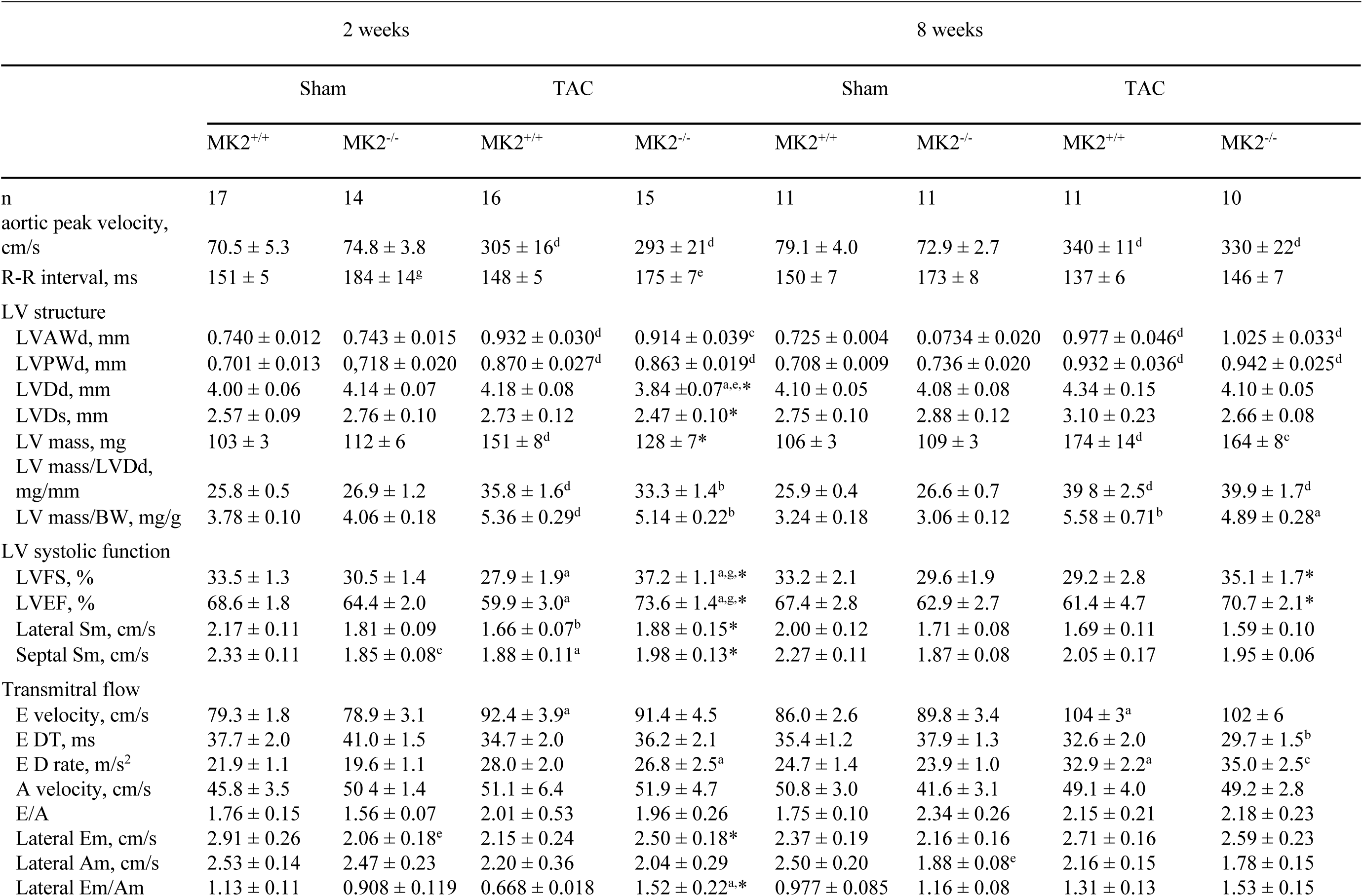
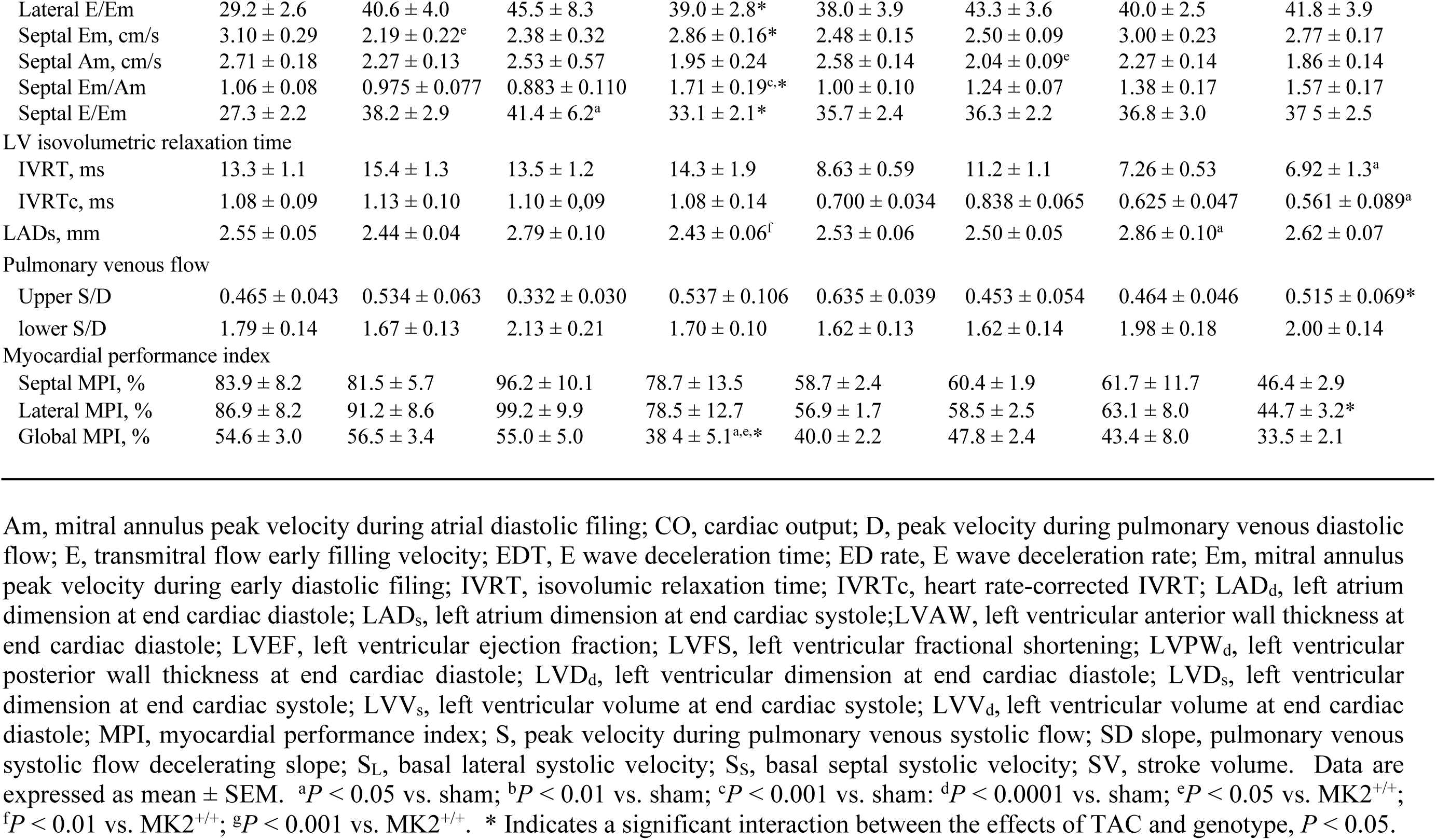
Echocardiographic parameters of MK2^+/+^ and MK2^−/−^ mice 2- and 8-weeks post-TAC.

Direct hemodynamic assessment by Millar catheter did not reveal major functional differences when between TAC-MK2^+/+^ and TAC-MK2^−/−^ mice; however, both 2 and 8 weeks post-TAC, MK2-deficient hearts showed a significant increase in the maximum -dP/dt relative to sham MK2^−/−^ mice whereas MK2^+/+^ mice did not (**Table 8**). Assessment of LV structure and function by echocardiographic imaging two-weeks post-TAC revealed reduced end-diastolic diameter (LVDd) in TAC MK2^−/−^ mice compared with both their sham counterparts (sham: 4.14 ± 0.07 mm, n = 14; TAC: 3.84 ± 0.07 mm, n = 15, *P* < 0.05) and TAC MK2^+/+^ mice (sham: 4.00 ± 0.06 mm, n = 17; TAC: 4.18 ± 0.08 mm, n = 16, *P* < 0.05). Eight weeks post-TAC these differences were no longer significant. Left ventricular anterior (LVAWd) and posterior (LVPWd) wall thickness was similarly increased in both TAC MK2^+/+^ and TAC MK2^−/−^ mice (ca. 20%, *P* < 0.001, **Table 9**). Left ventricular ejection fraction was reduced in MK2^+/+^ mice 2 weeks post-TAC (sham: 68.8 ± 1.8%, n = 17; TAC: 59.9 ± 3.0%, n = 16, *P* < 0.05) whereas it was increased in MK2^−/−^ mice (sham: 64.4 ± 2.0%, n = 14; TAC: 73.6 ± 1.4%, n = 15, *P* < 0.05) with LVEF being significantly greater in TAC MK2^−/−^ mice than TAC MK2^+/+^ mice (*P* < 0.001). Although these differences were also observed 8 weeks post-TAC, they did not reach statistical significance. However, 2-way ANOVA indicated a significant interaction between the effects of TAC and genotype on LV diameter (LVDd, LVDs), LV mass, and LV function (LVEF, LVFS) 2-weeks post-TAC (**Table 9**). Specifically, whereas LV diameter increased and ejection fraction decreased inMK2^+/+^ mice the opposite was observed in MK2-deficient mice. Furthermore, a significant interaction between the effects of TAC and genotype on LVEF and LVFS was also observed 8-weeks post-TAC.

Systolic (Sm) and early diastolic (Em) tissue velocity at the basal segments of the interventricular septum and lateral LV wall was assessed by tissue doppler imaging. As mentioned above, both lateral- and septal-Sm were lower in 12-week-old MK2^−/−^ mice relative to MK2^+/+^ mice **(Table 2)**. Two weeks post-TAC, systolic tissue velocities in TAC MK2^−/−^ mice did not differ from those in sham MK2^−/−^ mice **(Table 9)**. In contrast, two weeks post-TAC, lateral- and septal-Sm were significantly lower in TAC MK2^+/+^ relative to sham MK2^+/+^ mice. However, no differences in systolic tissue velocity were observed eight weeks post-TAC **(Table 9)**. Two-way ANOVA indicated a significant interaction between the effects of TAC and genotype on both Sm and Em (as well as E/Em and Em/Am), in both septal and lateral segments, 2-weeks post-TAC.

Early (E) and late (A) transmitral flow velocities, as well as the E/A ratio, did not differ significantly between TAC MK2^+/+^ and TAC MK2^−/−^ mice either two- or eight-weeks post-TAC **(Table 9)**. The E wave deceleration time (E DT) and isovolumetric relaxation time (IVRTc) were shortened significantly, relative to sham, in TAC MK2^−/−^ mice but not in in TAC MK2^+/+^ mice eight weeks post-TAC **(Figure 9A,B)**. Although these changes suggest LV compliance was reduced in TAC MK2^−/−^ mice, E DT and IVRTc in TAC MK2^−/−^ mice did not differ significantly from TAC MK2^+/+^ mice. In addition, both septal and lateral E/Em ratios, which reflect left ventricular filling pressure, did not differ. Similarly, the ratio of peak systolic to diastolic pulmonary venous flow velocity, which would increase in response to reduced LV compliance, was similar in TAC MK2^+/+^ and TAC MK2^−/−^ mice both two- and eight-weeks post-TAC. Furthermore, left atrial diameter, which may increase in response to a chronic increase in LV filling pressure (Abhayaratna *et al.*, 2006), increased in TAC MK2^+/+^ but not TAC MK2^−/−^ mice **(Figure 9C)**. Finally, the myocardial performance index, which increases as cardiac performance is reduced, was reduced in TAC MK2^−/−^ mice relative to both TAC MK2^+/+^ and sham MK2^−/−^ mice two weeks post-TAC, although this difference no longer reached significance eight weeks post-TAC **(Figure 9D)**. Two-way ANOVA indicated a significant interaction between the effects of TAC and genotype on global MPI 2-weeks post-TAC and on lateral MPI 8-weeks post-TAC (**Table 9**). Hence, MK2-deficiency resulted in a delay in hypertrophy and the onset of the changes in left ventricular structure and function evoked by a chronic increase in afterload.

**Figure. 9.**
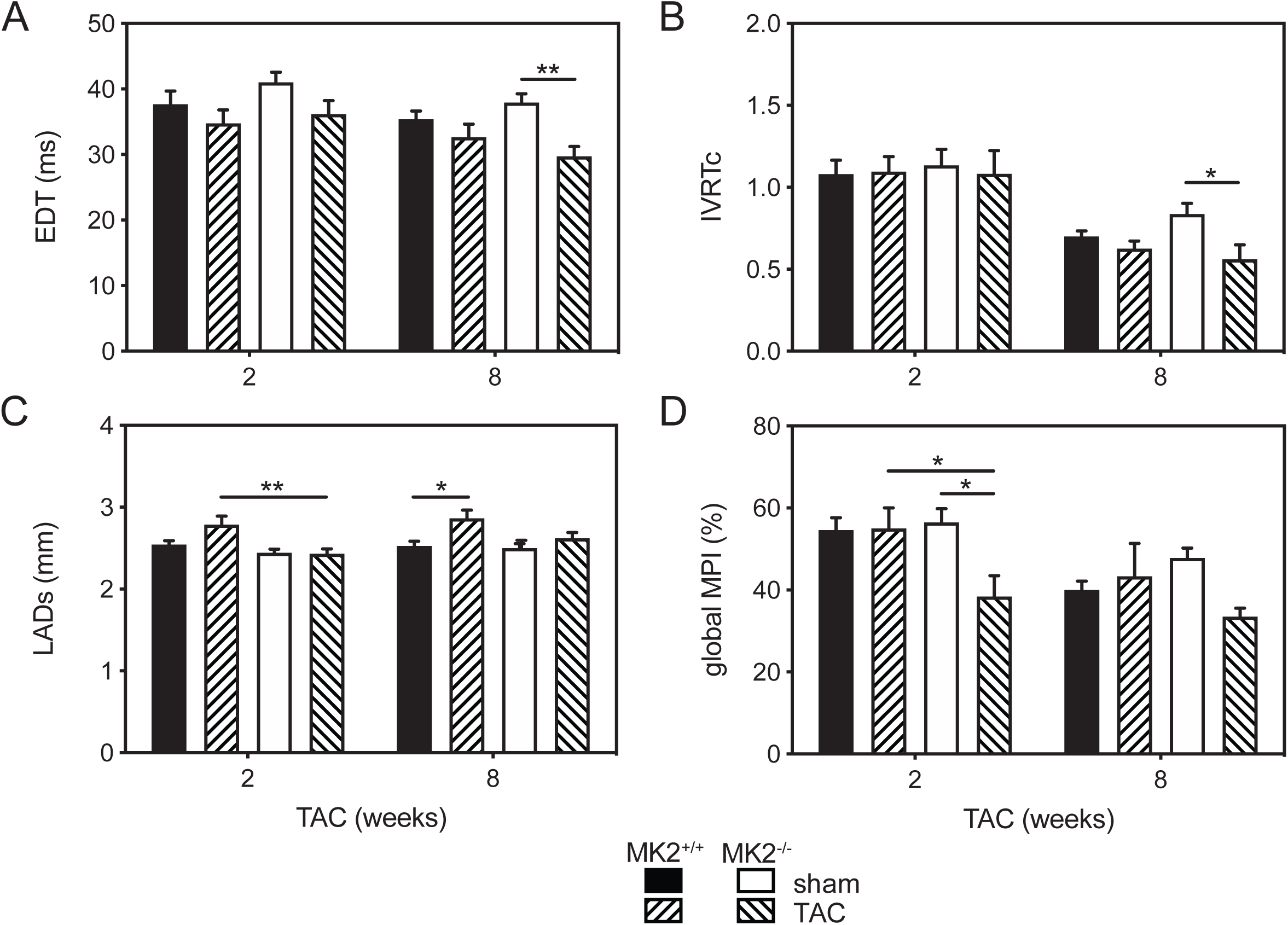
MK2-deficient hearts are not predisposed to further diastolic dysfunction when challenged with a chronic increase in afterload. Pressure overload was induced in MK2^−/−^ and wild-type littermate control (MK^+/+^) mice by constriction of the transverse aorta (TAC), and mice were euthanized 2 and 8 wk post-surgery. Sham-operated (sham) animals underwent the identical surgical procedure; however, the aorta was not constricted. **(A)** E wave deceleration time (EDT), **(A)** isovolumetric relaxation time corrected for differences in R-R interval (IVRTc), **(C)** left atrial diameter at systole (LADs), and **(D)** global myocardial performance index (MPI: note that higher values indicate poorer cardiac function). Data are presented as means ± SE; n = 6 – 17 mice. *, *P* < 0.05, \*\**P* < 0.01 by two-way ANOVA with Bonferroni’s multiple-comparison post-test.

Remodelling of the myocardium in response to a chronic increase in afterload involves both hypertrophy and increased interstitial fibrosis. We previously observed a modest fibrotic effect in mice on the mixed 129/Ola x C57BL genetic background (Nawaito *et al.*, 2017). Similarly, eight weeks post-TAC, Masson’s trichrome staining revealed very little increased interstitial fibrosis in TAC MK2^+/+^ and TAC MK2^−/−^ mice relative to their respective sham controls **(Figure 10)**.

**Figure 10.**
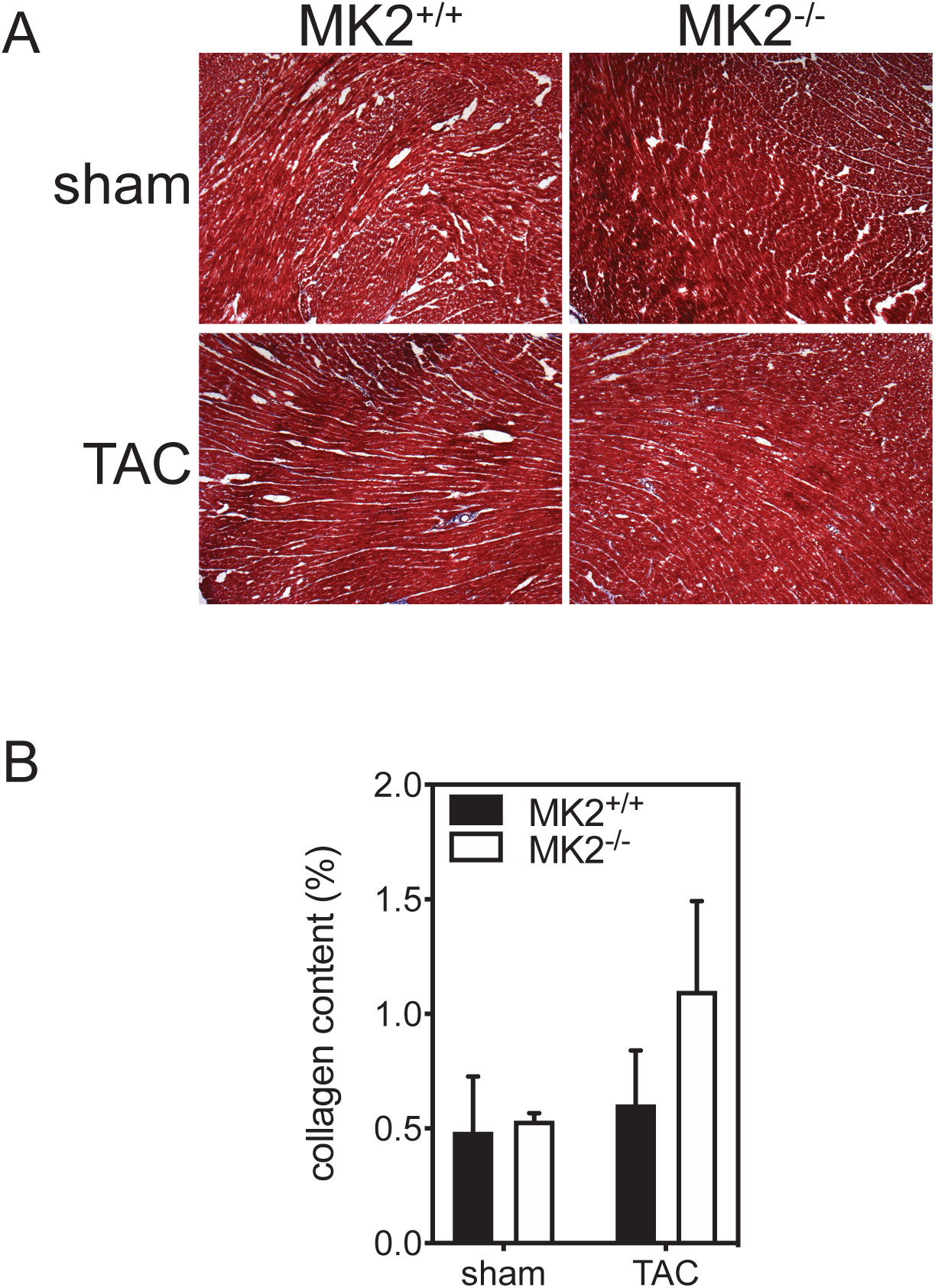
MK2 deficiency had no effect on interstitial fibrosis. **(A)** Transverse cryosections (8 mm) of the ventricular myocardium were stained with Masson’s trichrome 8 wk post-TAC. **(B)** collagen content was quantified by color segmentation. The original magnification was X40. Data are expressed as means ± SE; n = 6 - 8.

TAC activates the p38 pathway, with a peak activation occurring 3 days post-TAC (Esposito *et al.*, 2001; Dingar *et al.*, 2010). Furthermore, pharmacological inhibition of p38α/β potentiates ERK1/2 activation in cardiac myocytes (Boivin & Allen, 2012). Hence, we examined the effects of MK2-deficiency and TAC on phosphorylation of p38, MEK1/2, and ERK1/2 within their activation loops. Two weeks post-TAC, ERK1/2, MEK1/2 and p38 phosphorylation (normalized to total immunoreactivity) in TAC and sham hearts did not differ significantly. In each case, a modest, but not significant, reduction in phosphorylation was observed in the samples from MK2-deficient hearts (Data Not Shown). In addition, MK2 deficiency has been shown previously to destabilize p38α leading to a reduction p38α immunoreactivity (Kotlyarov *et al.*, 2002; Ruiz *et al.*, 2016). The abundance of p38α immunoreactivity was reduced in MK2-deficient hearts to a similar extent in sham and TAC hearts two weeks post-TAC (Data Not Shown).

## Discussion

Deletion of MK2 in mice had no detrimental impact on survival when examined up to 100 weeks-of-age. Interestingly, relative to age- and sex-matched littermates, MK2-deficient mice were hypothermic and had significantly lower body weight in later life. At twelve weeks of age, male MK2-deficient mice displayed prolonged RR intervals as well as signs of early diastolic dysfunction; however, diastolic dysfunction failed to develop with age. The amplitude and decay of calcium transients evoked by either field stimulation or caffeine were similar in ventricular myocytes isolated from adult male MK2^+/+^ and MK2^−/−^ mice whereas when perfused *ex vivo*, in working mode, hearts from MK2^+/+^ and MK2^−/−^ mice beat with similar frequency. Acquisition of ECG data in conscious mice by radio telemetry revealed P-R segment prolongation in MK2-deficient mice, suggesting autonomic regulation of heart function was altered in these mice. Finally, the LV hypertrophy induced by imposing a chronic increase in afterload was delayed, but not prevented, in MK2-deficient mice.

Echocardiographic imaging in 12-week-old male MK2^−/−^ mice revealed signs of reduced LV compliance, suggesting early diastolic dysfunction. However, MK2-deficient mice failed to develop diastolic dysfunction as they aged. Several factors are known to affect diastolic function, including preload (Fukuta & Little, 2008), aging (Nakou *et al.*, 2016), and heart rate (Esfandiari *et al.*, 2015). Although in theory the observed changes in diastolic function in MK2-deficient mice could be related to a decrease in heart rate, IVRT in MK2^−/−^ mice was significantly longer than MK2^+/+^ mice even after being corrected for heart rate. Among clinical strategies used to improve diastolic function, treatments, such as beta blockers, have been employed to both decrease heart rate and increase diastolic filling time (Ha & Oh, 2009). Hence, one possible explanation for the lack of progression to diastolic dysfunction in MK2^−/−^ mice could be a protective effect resulting from reduced heart rate.

Although bradycardic *in vivo*, when hearts from wild type and MK2-deficient mice were assessed using an *ex vivo* working heart preparation, heart rates did not differ, as shown previously (Ruiz *et al.*, 2016). This discrepancy can be explained by the non-negligible impact of differences in body temperature, neuronal, and/or hormonal regulation that would be absent in *ex vivo* heart preparations. Possible mechanisms involved in lowering the heart rate in MK2-deficient mice include changes in autonomic regulation of heart function: an enhanced parasympathetic regulation and/or a decreased sympathetic regulation could reduce heart rate (Gordan *et al.*, 2015). Consistent with this, when ECGs were acquired from conscious mice using radio telemetry, prolonged R-R intervals were observed in MK2-deficient mice along with prolongation of the P-R segment with no change in the P-wave-to-QRS-complex ratio. These observations are also consistent with there being alterations in autonomic regulation of heart function in the MK2-deficient mice. However, this hypothesis is based on basic functional observations, and further investigations will be required to better understand the role of MK2 in parasympathetic and sympathetic regulation of cardiac function, including the conduction velocity within the A-V node, and the G protein-coupled receptors involved in the catecholaminergic control of heart rate and contractility (Salazar *et al.*, 2007). Overall, the abovementioned results suggest a novel role for MK2 in regulating heart rate and diastolic function and highlight a new putative therapeutic target for heart diseases involving left ventricular diastolic dysfunction. Similarly, we recently reported that a deficiency in MK2 protects mice against diabetes-induced cardiac diastolic dysfunction (Ruiz *et al.*, 2016).

Mitochondria play a central role in cardiac energy production and transduction since mitochondrial oxidative phosphorylation is responsible for roughly 90% of the ATP production in the cardiomyocyte. A close link between mitochondrial failure and cardiac remodeling and dysfunction has been illustrated in various genetic models, particularly in mouse models lacking transcriptional coactivators and key drivers of mitochondrial biogenesis and function (Dorn *et al.*, 2015), such as PGC-1α (Arany *et al.*, 2006), PGC-1β (Riehle *et al.*, 2011), or both (Lai *et al.*, 2008). A similar link has been established in mouse (Faerber *et al.*, 2011) and in rat (Garnier *et al.*, 2003) models of heart failure as well as in the human failing heart following ischemic heart disease or dilated cardiomyopathy (Sebastiani *et al.*, 2007). In fact, heart failure is characterized by alterations in energy metabolism that includes changes in substrate utilization for energy production, particularly depressed fatty acid utilization, as well as changes in the expression of molecular regulators involved in the PGC-1α/PPARα axis (Neubauer, 2007; Lee & Kim, 2015). Therefore, targeting mitochondria in heart diseases, especially heart failure, through induction of modulators of mitochondrial biogenesis, for example, remains an active field of research aimed at identifying new therapeutic strategies (Bayeva *et al.*, 2013). In MK2-deficient mice, despite no significant changes in mitochondrial respiration rates and the activity of mitochondrial enzymes, other than succinate dehydrogenase, we observed an intriguing increase in the expression of key metabolic genes including PGC-1α and genes involved in fatty acid metabolism. MK2 has been shown to regulate the stability of mRNAs containing AU-rich elements (Neininger *et al.*, 2002; Ruiz *et al.*, 2018), which could explain at least part of the changes. Based on the role of depressed PGC-1α and fatty acid-related metabolic genes in mitochondrial dysfunction in heart failure, the fact that MK2-deficient mice exhibited enhanced expression of these latter genes and improved diastolic function, suggest that MK2 could be considered as an attractive target to improve mitochondrial function in heart diseases. Similar alterations in gene expression have been observed in skeletal muscle from mice where both MK2 and MK3 have been deleted (Scharf *et al.*, 2013). However, the implication of uncoupling protein, in heart failure is more controversial (Laskowski & Russell, 2008; Akhmedov *et al.*, 2015). Our results on UCP expression, including UCP3, which was increased in MK2-deficient mice, are more conflicting since some evidence supports a maladaptive effect of UCPs as key uncouplers of mitochondrial oxidative phosphorylation that decrease energetic efficiency (Murray *et al.*, 2008) while other studies pointed to an adaptive phenomenon of UCPs in response to lipid accumulation as exporters of fatty acids (Rial *et al.*, 2010) and by their ability to prevent ROS accumulation and cardiomyocyte apoptosis (Akhmedov *et al.*, 2015). Therefore, the significance of our findings needs to be further clarified and the metabolic profile of MK2-deficient hearts under conditions of stress assessed.

Among mechanisms involved in mitochondrial pathogenesis is the opening of the mitochondrial permeability transition pore (MPTP), which results in uncoupling of the mitochondrial respiratory chain, impaired ATP synthesis, and eventually mitochondrial swelling and cell death (Kwong & Molkentin, 2015; Perez & Quintanilla, 2017). The exact molecular identity of this complex has not yet been well defined and is the subject of intense debate although the molecular structure of MPTP includes the volt-dependent anion channel (VDAC), the adenine nucleotide translocator (ANT), and cyclophilin D, among other possible components that include the F1F0 ATP synthase for the pore-forming component (Bernardi & Di Lisa, 2015; Kwong & Molkentin, 2015). Nevertheless, increased or sustained MPTP opening subsequent to ROS and/or Ca^2+^ overload is involved in mitochondrial membrane permeabilization, which in turn results in mitochondrial respiratory chain uncoupling as well as impaired oxidative phosphorylation, blunted ATP production, and ultimately mitochondrial swelling and cell death (Kinnally *et al.*, 2011; Perez & Quintanilla, 2017). There is increasing evidence supporting a role for PTP opening in cardiac diseases. In ischemia-reperfusion injury, prolonged MPTP opening has been described as a key contributor to post-ischemia/reperfusion-mediated cardiomyocyte death (Di Lisa *et al.*, 2001). Similarly, in response to volume overload, mitochondrial vulnerability to PTP opening increased during compensated ventricular hypertrophy (Marcil *et al.*, 2006; Matas *et al.*, 2009). In rats subjected to coronary artery ligation, MPTP opening was prolonged during myocardial infarction (Javadov *et al.*, 2005). Finally, MPTP opening appears to persist in failing hearts following chronic pacing or post-infarction remodeling in dogs (Sharov *et al.*, 2007) as well as in rats following decompensated hypertrophy secondary to TAC-induced increased afterload (Dabkowski *et al.*, 2013). Therefore, targeting MPTP opening is now considered to be a promising therapeutic target for the treatment of cardiac diseases and new strategies aimed at limiting MPTP opening are required to overcome the existing challenges that limit the development of specific MPTP inhibitors (Javadov *et al.*, 2009;Bernardi & Di, 2015). Our results show that inhibiting MK2 signaling reduced the Ca^2+^-sensitivity for MPTP opening; hence, MK2 or its down-stream targets may warrant consideration in the development novel cardioprotective therapies.

When the effects of a chronic increase in afterload on cardiac remodeling were examined in MK2-deficient mice, the findings were consistent with those of Streicher *et al* (Streicher *et al.*, 2010). By inducing the overexpression of a constitutively-active form of MKK3 (MKK3E) in a cardiomyocyte-specific manner, these authors showed that MK2 participates in the p38-mediated pathological cardiac remodeling and contractile dysfunction. In addition, a deficiency of MK2 attenuates ventricular hypertrophy and improves contractility, despite no difference in the expression molecular markers of remodeling. Similarly, our results show that following TAC, MK2-deficient mice showed reduced hypertrophy and improved contractility, despite no difference in the expression of hypertrophic markers. More specifically, two-weeks after TAC, MK2-deficient mice showed improved: (i) left ventricular systolic function as well as (ii) left ventricular diastolic properties. These findings do not entirely concur with the findings of Streicher *et al.* who further showed that knocking out MK2 in MKK3E-overexpressing hearts does not rescue myocardial stiffness (Streicher *et al.*, 2010). This discrepancy can be explained by the different model used: chronic activation of p38 MAPK versus TAC-induced increased afterload. Being a downstream target of p38α/β, MK2 could be contributing to the pathogenic responses of p38 activation by either modulating total p38α protein levels or mediated downstream signaling. In fact, in MK2-deficient mice, the amount of p38α is significantly reduced (Ruiz *et al.*, 2016), since MK2 stabilizes p38α via a direct interaction with its C-terminal (Kotlyarov *et al.*, 2002). Alternatively, downstream targets of MK2, such as hsp25/27, may mediate molecular events induced by MK2 activation. Overexpression of a non-phosphorylatable hsp27 mutant improves cardiac performance (Clements *et al.*, 2011). However, it is noteworthy that the benefits of MK2 loss observed 2 weeks post-TAC were less pronounced when examined 8 weeks post-TAC albeit hearts from MK2^−/−^ mice after 8 wk-TAC showed a better compliance. These differences suggest that rather than abrogating pressure overload-induced contractile dysfunction, the absence of MK2 delays the pathogenic response.

### Potential limitations

One limitation of the present study is the animal model. The MK2-deficient mice are pan knockouts rather than cardiomyocyte-specific. Hence, the involvement of body temperature, peripheral mechanisms, or systemic mediators in the observed cardiac phenotype cannot be excluded. In addition, although the overall aim of this study was to perform a comprehensive analysis of the role of MK2 in regulating basal cardiac function and its implication in pathological remodeling, our results do not shed light on the molecular mechanisms by which MK2 regulates cardiac function in health and disease, thereby representing another limitation.

### Conclusions

MK2-deficient mice showed no adverse effects on myocardial performance over time. Interestingly, autonomic regulation of cardiac function appears to be altered in the absence of MK2. In addition, MK2-deficient mice showed changes in the expression of some metabolic genes related to mitochondrial biogenesis and fatty acid metabolism and, despite no alterations in mitochondrial respiration rates. Furthermore, the calcium-sensitivity for mPTP opening was reduced, suggesting a cardioprotective effect of MK2 loss. Finally, in response to a chronic increase in afterload, left ventricular hypertrophy was delayed, but not reduced or prevented. Altogether these findings support a novel role for MK2 in the heart that merits further investigation, especially in regard to its possible role in autonomic regulation of heart function, the regulation of metabolic gene expression, and mPTP opening as these may represent interesting cardioprotective roles for inhibitors of MK2. In addition, consistent with previous work using a constitutive form of MK2 deficiency, we confirm that MK2 is implicated in the pathological processes leading to cardiac hypertrophy and ventricular dysfunction.

## Abbreviations

ANF: atrial natriuretic factor
BW: body weight
CS: citrate synthase
DMSO: dimethyl sulfoxide
DTT: dithiothreitol
E: early diastolic transmitral filling velocity
EDT: early diastolic transmitral filling deceleration time
Em: peak early diastolic tissue velocity
HW: heart weight
HSL: hormone-sensitive lipase
ICDH: isocitrate dehydrogenase
IVRTc: rate-corrected isovolumic relaxation time
LV: left ventricle
LVAWd: left ventricle anterior wall end-diastolic thickness
LVPWd: left ventricle posterior wall end-diastolic thickness
LVEDD: left ventricular end-diastolic diameter
LVESD: left ventricular end-systolic diameter
FS: fractional shortening
LVEF: left ventricular ejection fraction
MAPK: mitogen-activated protein kinase
MCAD: medium-chain acyl coenzyme A (acyl-CoA) dehydrogenase
MHCβ: myosin heavy chain β
MK2: MAPK-activated protein kinase-2
MK3: MAPK-activated protein kinase-3
MKK3: MAPK kinase 3
MKK6: MAPK kinase 6
β-MHC: β-myosin heavy chain
MPI: myocardial performance index
PAGE: polyacrylamide gel electrophoresis
PGC-1α: peroxisome proliferator-activated receptor γ coactivator 1-α
PKA: protein kinase A
PMSF: phenylmethylsulfonyl fluoride
PRAK: p38-regulated/activated protein kinase
SSM: subsarcolemmal mitochondria
TAC: transverse aortic constriction
TDI: tissue Doppler imaging
TX-100: Triton X-100
UCP3: uncoupling protein 3

## Additional information

### Author contributions

MR, MK, DD, GV, JT. RJK, YS, BH, AN, BGA conceived and designed the experiments. MR, MK, DD, GV, JT. RJK, YS, BH, AN, SAN, and PS performed the experiments and analyzed the data. MR, MK, and BGA assembled and interpreted all the data. MG provided the mouse model. MR, MK and BGA wrote and critically reviewed the manuscript. All authors have approved the final version of the current manuscript.

## Acknowledgements

We thank Ms. Karine Bouthillier and Dr. Robert Parent for animal care and breeding as well as Dr Christine Des Rosiers for her thoughtful comments.

## References

Abhayaratna WP, Seward JB, Appleton CP, Douglas PS, Oh JK, Tajik AJ & Tsang TS. (2006). Left atrial size: physiologic determinants and clinical applications. J Am Coll Cardiol 47, 2357–2363.

Akashi YJ, Springer J, Lainscak M & Anker SD. (2007). Atrial natriuretic peptide and related peptides. Clin Chem Lab Med 45, 1259–1267.

Akhmedov AT, Rybin V & Marin-Garcia J. (2015). Mitochondrial oxidative metabolism and uncoupling proteins in the failing heart. Heart Fail Rev 20, 227–249.

Arabacilar P & Marber M. (2015). The case for inhibiting p38 mitogen-activated protein kinase in heart failure. Front Pharmacol 6, 102.

Arany Z, Novikov M, Chin S, Ma Y, Rosenzweig A & Spiegelman BM. (2006). Transverse aortic constriction leads to accelerated heart failure in mice lacking PPAR-gamma coactivator 1alpha. Proc Natl Acad Sci U S A 103, 10086–10091.

Arthur JS & Ley SC. (2013). Mitogen-activated protein kinases in innate immunity. Nat Rev Immunol 13, 679–692.

Bayeva M, Gheorghiade M & Ardehali H. (2013). Mitochondria as a therapeutic target in heart failure. J Am Coll Cardiol 61, 599–610.

Bellahcene M, Jacquet S, Cao XB, Tanno M, Haworth RS, Layland J, Kabir AM, Gaestel M, Davis RJ, Flavell RA, Shah AM, Avkiran M & Marber MS. (2006). Activation of p38 mitogen-activated protein kinase contributes to the early cardiodepressant action of tumor necrosis factor. J Am Coll Cardiol 48, 545–555.

Bernardi P & Di Lisa F. (2015). The mitochondrial permeability transition pore: molecular nature and role as a target in cardioprotection. J Mol Cell Cardiol 78, 100–106.

Boivin B & Allen BG. (2012). p38 MAP kinase attenuates phorbol ester-induced ERK MAP kinase activation in adult cardaic ventricular myocytes. Curr Topics Biochem Res 14, 57–63.

Braz JC, Bueno OF, Liang Q, Wilkins BJ, Dai YS, Parsons S, Braunwart J, Glascock BJ, Klevitsky R, Kimball TF, Hewett TE & Molkentin JD. (2003). Targeted inhibition of p38 MAPK promotes hypertrophic cardiomyopathy through upregulation of calcineurin-NFAT signaling. J Clin Invest 111, 1475–1486.

Brouillette J, Grandy SA, Jolicoeur P & Fiset C. (2007). Cardiac repolarization is prolonged in CD4C/HIV transgenic mice. J Mol Cell Cardiol 43, 159–167.

Cargnello M & Roux PP. (2011). Activation and function of the MAPKs and their substrates, the MAPK-activated protein kinases. Microbiol Mol Biol Rev 75, 50–83.

Chahine MN, Mioulane M, Sikkel MB, O’Gara P, Dos Remedios CG, Pierce GN, Lyon AR, Foldes G & Harding SE. (2015). Nuclear pore rearrangements and nuclear trafficking in cardiomyocytes from rat and human failing hearts. Cardiovasc Res 105, 31–43.

Cheung PC, Campbell DG, Nebreda AR & Cohen P. (2003). Feedback control of the protein kinase TAK1 by SAPK2a/p38α. EMBO J 22, 5793–5805.

Clements RT, Feng J, Cordeiro B, Bianchi C & Sellke FW. (2011). p38 MAPK-dependent small HSP27 and αB-crystallin phosphorylation in regulation of myocardial function following cardioplegic arrest. Am J Physiol Heart Circ Physiol 300, H1669–1677.

Connern CP & Halestrap AP. (1996). Chaotropic agents and increased matrix volume enhance binding of mitochondrial cyclophilin to the inner mitochondrial membrane and sensitize the mitochondrial permeability transition to [Ca^2+^]. Biochemistry 35, 8172–8180.

Coulthard LR, White DE, Jones DL, McDermott MF & Burchill SA. (2009). p38^MAPK^: stress responses from molecular mechanisms to therapeutics. Trends Mol Med 15, 369–379.

Dabkowski ER, O’Connell KA, Xu W, Ribeiro RF, Jr., Hecker PA, Shekar KC, Daneault C, Des Rosiers C & Stanley WC. (2013). Docosahexaenoic acid supplementation alters key properties of cardiac mitochondria and modestly attenuates development of left ventricular dysfunction in pressure overload-induced heart failure. Cardiovasc Drugs Ther 27, 499–510.

Depre C, Shipley GL, Chen W, Han Q, Doenst T, Moore ML, Stepkowski S, Davies PJ & Taegtmeyer H. (1998). Unloaded heart in vivo replicates fetal gene expression of cardiac hypertrophy. Nat Med 4, 1269–1275.

Di Lisa F, Menabo R, Canton M, Barile M & Bernardi P. (2001). Opening of the mitochondrial permeability transition pore causes depletion of mitochondrial and cytosolic NAD^+^ and is a causative event in the death of myocytes in postischemic reperfusion of the heart. J Biol Chem 276, 2571–2575.

Dingar D, Merlen C, Grandy S, Gillis MA, Villeneuve LR, Mamarbachi AM, Fiset C & Allen BG. (2010). Effect of pressure overload-induced hypertrophy on the expression and localization of p38 MAP kinase isoforms in the mouse heart. Cell Signal 22, 1634–1644.

Dorn GW, 2nd, Vega RB & Kelly DP. (2015). Mitochondrial biogenesis and dynamics in the developing and diseased heart. Genes Dev 29, 1981–1991.

Duda MK, O’Shea KM, Tintinu A, Xu W, Khairallah RJ, Barrows BR, Chess DJ, Azimzadeh AM, Harris WS, Sharov VG, Sabbah HN & Stanley WC. (2009). Fish oil, but not flaxseed oil, decreases inflammation and prevents pressure overload-induced cardiac dysfunction. Cardiovasc Res 81, 319–327.

Dulos J, Wijnands FP, van den Hurk-van Alebeek JA, van Vugt MJ, Rullmann JA, Schot JJ, de Groot MW, Wagenaars JL, van Ravestein-van Os R, Smets RL, Vink PM, Hofstra CL, Nelissen RL & van Eenennaam H. (2013). p38 inhibition and not MK2 inhibition enhances the secretion of chemokines from TNF-alpha activated rheumatoid arthritis fibroblast-like synoviocytes. Clin Exp Rheumatol 31, 515–525.

El Khoury N, Mathieu S, Marger L, Ross J, El Gebeily G, Ethier N & Fiset C. (2013). Upregulation of the hyperpolarization-activated current increases pacemaker activity of the sinoatrial node and heart rate during pregnancy in mice. Circulation 127, 2009–2020.

Esfandiari S, Fuchs F, Wainstein RV, Chelvanathan A, Mitoff P, Sasson Z & Mak S. (2015). Heart rate-dependent left ventricular diastolic function in patients with and without heart failure. J Card Fail 21, 68–75.

Esposito G, Prasad SVN, Rapacciuolo A, Mao L, Koch WJ & Rockman HA. (2001). Cardiac overexpression of a Gq inhibitor blocks induction of extracellular signal–regulated kinase and c-Jun NH_2_-terminal kinase activity in in vivo pressure overload. Circulation 103, 1453–1458.

Faerber G, Barreto-Perreia F, Schoepe M, Gilsbach R, Schrepper A, Schwarzer M, Mohr FW, Hein L & Doenst T. (2011). Induction of heart failure by minimally invasive aortic constriction in mice: reduced peroxisome proliferator-activated receptor gamma coactivator levels and mitochondrial dysfunction. J Thorac Cardiovasc Surg 141, 492–500.

Fiore M, Forli S & Manetti F. (2016). Targeting mitogen-activated protein kinase-activated protein kinase 2 (MAPKAPK2, MK2): medicinal chemistry efforts to lead small molecule inhibitors to clinical trials. J Med Chem 59, 3609–3634.

Frey N, Katus HA, Olson EN & Hill JA. (2004). Hypertrophy of the heart: a new therapeutic target? Circulation 109, 1580–1589.

Frey N, Luedde M & Katus HA. (2011). Mechanisms of disease: hypertrophic cardiomyopathy. Nat Rev Cardiol 9, 91–100.

Fukuta H & Little WC. (2008). The cardiac cycle and the physiologic basis of left ventricular contraction, ejection, relaxation, and filling. Heart Fail Clin 4, 1–11.

Garnier A, Fortin D, Delomenie C, Momken I, Veksler V & Ventura-Clapier R. (2003). Depressed mitochondrial transcription factors and oxidative capacity in rat failing cardiac and skeletal muscles. J Physiol 551, 491–501.

Gelinas R, Labarthe F, Bouchard B, Mc Duff J, Charron G, Young ME & Des Rosiers C. (2008). Alterations in carbohydrate metabolism and its regulation in PPARalpha null mouse hearts. Am J Physiol Heart Circ Physiol 294, H1571–H1580.

Gonzalez-Teran B, Lopez JA, Rodriguez E, Leiva L, Martinez-Martinez S, Bernal JA, Jimenez-Borreguero LJ, Redondo JM, Vazquez J & Sabio G. (2016). p38γ and δ promote heart hypertrophy by targeting the mTOR-inhibitory protein DEPTOR for degradation. Nat Commun 7, 10477.

Gordan R, Gwathmey JK & Xie LH. (2015). Autonomic and endocrine control of cardiovascular function. World J Cardiol 7, 204–214.

Ha JW & Oh JK. (2009). Therapeutic strategies for diastolic dysfunction: a clinical perspective. J Cardiovasc Ultrasound 17, 86–95.

Kaul S, Tei C, Hopkins JM & Shah PM. (1984). Assessment of right ventricular function using two-dimensional echocardiography. Am Heart J 107, 526–531.

Khairallah M, Labarthe F, Bouchard B, Danialou G, Petrof BJ & Des Rosiers C. (2004). Profiling substrate fluxes in the isolated working mouse heart using 13C-labeled substrates: focusing on the origin and fate of pyruvate and citrate carbons. Am J Physiol Heart Circ Physiol 286, H1461–H1470.

Khairallah RJ, O’Shea KM, Brown BH, Khanna N, Des Rosiers C & Stanley WC. (2010a). Treatment with docosahexaenoic acid, but not eicosapentaenoic acid, delays Ca^2+^-induced mitochondria permeability transition in normal and hypertrophied myocardium. J Pharmacol Exp Ther 335, 155–162.

Khairallah RJ, Sparagna GC, Khanna N, O’Shea KM, Hecker PA, Kristian T, Fiskum G, Des Rosiers C, Polster BM & Stanley WC. (2010b). Dietary supplementation with docosahexaenoic acid, but not eicosapentaenoic acid, dramatically alters cardiac mitochondrial phospholipid fatty acid composition and prevents permeability transition. Biochim Biophys Acta 1797, 1555–1562.

Kinnally KW, Peixoto PM, Ryu SY & Dejean LM. (2011). Is mPTP the gatekeeper for necrosis, apoptosis, or both? Biochim Biophys Acta 1813, 616–622.

Kong P, Christia P & Frangogiannis NG. (2014). The pathogenesis of cardiac fibrosis. Cell Mol Life Sci 71, 549–574.

Kotlyarov A, Neininger A, Schubert C, Eckert R, Birchmeier C, Volk HD & Gaestel M. (1999). MAPKAP kinase 2 is essential for LPS-induced TNF-α biosynthesis. Nat Cell Biol 1, 94–97.

Kotlyarov A, Yannoni Y, Fritz S, Laaβ K, Telliez JB, Pitman D, Lin LL & Gaestel M. (2002). Distinct cellular functions of MK2. Mol Cell Biol 22, 4827–4835.

Kwong JQ & Molkentin JD. (2015). Physiological and pathological roles of the mitochondrial permeability transition pore in the heart. Cell Metab 21, 206–214.

Lai L, Leone TC, Zechner C, Schaeffer PJ, Kelly SM, Flanagan DP, Medeiros DM, Kovacs A & Kelly DP. (2008). Transcriptional coactivators PGC-1alpha and PGC-lbeta control overlapping programs required for perinatal maturation of the heart. Genes Dev 22, 1948–1961.

Laskowski KR & Russell RR, 3rd. (2008). Uncoupling proteins in heart failure. Curr Heart Fail Rep 5, 75–79.

Lee WS & Kim J. (2015). Peroxisome proliferator-activated receptors and the heart: lessons from the past and future directions. PPAR Res 2015, 271983.

Lemke LE, Bloem LJ, Fouts R, Esterman M, Sandusky G & Vlahos CJ. (2001). Decreased p38 MAPK activity in end-stage failing human myocardium: p38 MAPK alpha is the predominant isoform expressed in human heart. J Mol Cell Cardiol 33, 1527–1540.

Marber MS, Rose B & Wang Y. (2011). The p38 mitogen-activated protein kinase pathway--a potential target for intervention in infarction, hypertrophy, and heart failure. J Mol Cell Cardiol 51, 485–490.

Marcil M, Ascah A, Matas J, Belanger S, Deschepper CF & Burelle Y. (2006). Compensated volume overload increases the vulnerability of heart mitochondria without affecting their functions in the absence of stress. J Mol Cell Cardiol 41, 998–1009.

Martin ED, Bassi R & Marber MS. (2015). p38 MAPK in cardioprotection - are we there yet? Br J Pharmacol 172, 2101–2113.

Matas J, Young NT, Bourcier-Lucas C, Ascah A, Marcil M, Deschepper CF & Burelle Y. (2009). Increased expression and intramitochondrial translocation of cyclophilin-D associates with increased vulnerability of the permeability transition pore to stress-induced opening during compensated ventricular hypertrophy. J Mol Cell Cardiol 46, 420–430.

Merlet N, Busseuil D, Mihalache-Avram T, Mecteau M, Shi Y, Nachar W, Brand G, Brodeur MR, Charpentier D, Rhainds D, Sy G, Schwendeman A, Lalwani N, Dasseux JL, Rheaume E & Tardif JC. (2016). HDL mimetic peptide CER-522 treatment regresses left ventricular diastolic dysfunction in cholesterol-fed rabbits. Int J Cardiol 215, 364–371.

Mitchell GF, Jeron A & Koren G. (1998). Measurement of heart rate and Q-T interval in the conscious mouse. Am J Physiol 274, H747–H751.

Murray AJ, Cole MA, Lygate CA, Carr CA, Stuckey DJ, Little SE, Neubauer S & Clarke K. (2008). Increased mitochondrial uncoupling proteins, respiratory uncoupling and decreased efficiency in the chronically infarcted rat heart. J Mol Cell Cardiol 44, 694–700.

Nakou ES, Parthenakis FI, Kallergis EM, Marketou ME, Nakos KS & Vardas PE. (2016). Healthy aging and myocardium: A complicated process with various effects in cardiac structure and physiology. Int J Cardiol 209, 167–175.

Nawaito SA, Dingar D, Sahadevan P, Hussein B, Sahmi F, Shi Y, Gillis MA, Gaestel M, Tardif JC & Allen BG. (2017). MK5 haplodeficiency attenuates hypertrophy and preserves diastolic function during remodeling induced by chronic pressure overload in the mouse heart. Am J Physiol Heart Circ Physiol 313, H46–H58.

Nawaito SA, Sahadevan P, Sahmi F, Gaestel M, Calderone A & Allen BG. (2019). Transcript levels for extracellular matrix proteins are altered in MK5-deficient cardiac ventricular fibroblasts. J Mol Cell Cardiol 132, 164–177.

Neininger A, Kontoyiannis D, Kotlyarov A, Winzen R, Eckert R, Volk HD, Holtmann H, Kollias G & Gaestel M. (2002). MK2 targets AU-rich elements and regulates biosynthesis of tumor necrosis factor and interleukin-6 independently at different post-transcriptional levels. J Biol Chem 277, 3065–3068.

Nemoto S, Sheng Z & Lin A. (1998). Opposing effects of Jun kinase and p38 mitogen-activated protein kinases on cardiomyocyte hypertrophy. Mol Cell Biol 18, 3518–3526.

Neubauer S. (2007). The failing heart--an engine out of fuel. N Engl J Med 356, 1140–1151.

Nishida K, Yamaguchi O, Hirotani S, Hikoso S, Higuchi Y, Watanabe T, Takeda T, Osuka S, Morita T, Kondoh G, Uno Y, Kashiwase K, Taniike M, Nakai A, Matsumura Y, Miyazaki J, Sudo T, Hongo K, Kusakari Y, Kurihara S, Chien KR, Takeda J, Hori M & Otsu K. (2004). p38α mitogen-activated protein kinase plays a critical role in cardiomyocyte survival but not in cardiac hypertrophic growth in response to pressure overload. Mol Cell Biol 24, 10611–10620.

O’Shea KM, Khairallah RJ, Sparagna GC, Xu W, Hecker PA, Robillard-Frayne I, Des Rosiers C, Kristian T, Murphy RC, Fiskum G & Stanley WC. (2009). Dietary omega-3 fatty acids alter cardiac mitochondrial phospholipid composition and delay Ca2+-induced permeability transition. J Mol Cell Cardiol 47, 819–827.

Perez MJ & Quintanilla RA. (2017). Development or disease: duality of the mitochondrial permeability transition pore. Dev Biol 426, 1–7.

Rial E, Rodriguez-Sanchez L, Gallardo-Vara E, Zaragoza P, Moyano E & Gonzalez-Barroso MM. (2010). Lipotoxicity, fatty acid uncoupling and mitochondrial carrier function. Biochim Biophys Acta 1797, 800–806.

Riehle C, Wende AR, Zaha VG, Pires KM, Wayment B, Olsen C, Bugger H, Buchanan J, Wang X, Moreira AB, Doenst T, Medina-Gomez G, Litwin SE, Lelliott CJ, Vidal-Puig A & Abel ED. (2011). PGC-1β deficiency accelerates the transition to heart failure in pressure overload hypertrophy. Circ Res 109, 783–793.

Rivard K, Grandy SA, Douillette A, Paradis P, Nemer M, Allen BG & Fiset C. (2011). Overexpression of type 1 angiotensin II receptors impairs excitation-contraction coupling in the mouse heart. Am J Physiol Heart Circ Physiol 301, H2018–H2027.

Ruiz M, Coderre L, Allen BG & Des Rosiers C. (2018). Protecting the heart through MK2 modulation, toward a role in diabetic cardiomyopathy and lipid metabolism. Biochim Biophys Acta 1864, 1914–1922.

Ruiz M, Coderre L, Lachance D, Houde V, Martel C, Thompson Legault J, Gillis MA, Bouchard B, Daneault C, Carpentier AC, Gaestel M, Allen BG & Des Rosiers C. (2016). MK2 deletion in mice prevents diabetes-induced perturbations in lipid metabolism and cardiac dysfunction. Diabetes 65, 381–392.

Ruiz M, Gelinas R, Vaillant F, Lauzier B & Des Rosiers C. (2015). Metabolic tracing using stable isotope-labeled substrates and mass spectrometry in the perfused mouse heart. Methods Enzymol 561, 107–147.

Sadoshima J & Izumo S. (1997). The cellular and molecular response of cardiac myocytes to mechanical stress. Annu Rev Physiol 59, 551–571.

Salazar NC, Chen J & Rockman HA. (2007). Cardiac GPCRs: GPCR signaling in healthy and failing hearts. Biochim Biophys Acta 1768, 1006–1018.

Scharf M, Neef S, Freund R, Geers-Knörr C, Franz-Wachtel M, Brandis A, Krone D, Schneider H, Groos S, Menon MB, Chang KC, Kraft T, Meissner JD, Boheler KR, Maier LS, Gaestel M & Scheibe RJ. (2013). Mitogen-activated protein kinase-activated protein kinases 2 and 3 regulate SERCA2a expression and fiber type composition to modulate skeletal muscle and cardiomyocyte function. Mol Cell Biol 33, 2586–2602.

Sebastiani M, Giordano C, Nediani C, Travaglini C, Borchi E, Zani M, Feccia M, Mancini M, Petrozza V, Cossarizza A, Gallo P, Taylor RW & d’Amati G. (2007). Induction of mitochondrial biogenesis is a maladaptive mechanism in mitochondrial cardiomyopathies. J Am Coll Cardiol 50, 1362–1369.

Shaffer F & Ginsberg JP. (2017). An overview of heart rate variability metrics and norms. Front Public Health 5, 258.

Shah AM & Mann DL. (2011). In search of new therapeutic targets and strategies for heart failure: recent advances in basic science. Lancet 378, 704–712.

Sharov VG, Todor A, Khanal S, Imai M & Sabbah HN. (2007). Cyclosporine A attenuates mitochondrial permeability transition and improves mitochondrial respiratory function in cardiomyocytes isolated from dogs with heart failure. J Mol Cell Cardiol 42, 150–158.

Shiroto K, Otani H, Yamamoto F, Huang CK, Maulik N & Das DK. (2005). MK2^−/−^ gene knockout mouse hearts carry anti-apoptotic signal and are resistant to ischemia reperfusion injury. J Mol Cell Cardiol 38, 93–97.

Streicher JM, Ren S, Herschman H & Wang Y. (2010). MAPK-activated protein kinase-2 in cardiac hypertrophy and cyclooxygenase-2 regulation in heart. Circ Res 106, 1434–1443.

Takeishi Y, Huang Q, Abe J, Che W, Lee JD, Kawakatsu H, Hoit BD, Berk BC & Walsh RA. (2002). Activation of mitogen-activated protein kinases and p90 ribosomal S6 kinase in failing human hearts with dilated cardiomyopathy. Cardiovasc Res 53, 131–137.

Trempolec N, Dave-Coll N & Nebreda AR. (2013a). SnapShot: p38 MAPK signaling. Cell 152, 656–656.

Trempolec N, Dave-Coll N & Nebreda AR. (2013b). SnapShot: p38 MAPK substrates. Cell 152, 924–924.

Verweij N, Mateo Leach I, van den Boogaard M, van Veldhuisen DJ, Christoffels VM, LifeLines Cohort S, Hillege HL, van Gilst WH, Barnett P, de Boer RA & van der Harst P. (2014). Genetic determinants of P wave duration and PR segment. Circ Cardiovasc Genet 7, 475–481.

Wang Y, Huang S, Sah VP, Ross J, Jr., Brown JH, Han J & Chien KR. (1998). Cardiac muscle cell hypertrophy and apoptosis induced by distinct members of the p38 mitogen-activated protein kinase family. J Biol Chem 273, 2161–2168.

Xu L, Yates CC, Lockyer P, Xie L, Bevilacqua A, He J, Lander C, Patterson C & Willis M. (2014). MMI-0100 inhibits cardiac fibrosis in myocardial infarction by direct actions on cardiomyocytes and fibroblasts via MK2 inhibition. J Mol Cell Cardiol 77, 86–101.

Yokota T & Wang Y. (2016). p38 MAP kinases in the heart. Gene 575, 369–376.

Zhang S, Weinheimer C, Courtois M, Kovacs A, Zhang CE, Cheng AM, Wang Y & Muslin AJ. (2003). The role of the Grb2-p38 MAPK signaling pathway in cardiac hypertrophy and fibrosis. J Clin Invest 111, 833–841.

